# An MRI-Derived Neuroanatomical Atlas of the Fischer 344 Rat Brain

**DOI:** 10.1101/743583

**Authors:** Dana Goerzen, Caitlin Fowler, Gabriel A. Devenyi, Jurgen Germann, Dan Madularu, M. Mallar Chakravarty, Jamie Near

## Abstract

This paper reports the development of a high-resolution 3-D MRI atlas of the Fischer 344 adult rat brain. The atlas is a 60 μm isotropic image volume composed of 256 coronal slices with 71 manually delineated structures and substructures. The atlas was developed using Pydpiper image registration pipeline to create an average brain image of 41 four-month-old male and female Fischer 344 rats. Slices in the average brain image were then manually segmented, individually and bilaterally, on the basis of image contrast in conjunction with Paxinos and Watson’s (2007) stereotaxic rat brain atlas. Summary statistics (mean and standard deviation of regional volumes) are reported for each brain region across the sample used to generate the atlas, and a statistical comparison of a chosen subset of regional brain volumes between male and female rats is presented. On average, the coefficient of variation of regional brain volumes across all rats in our sample was 4%, with no individual brain region having a coefficient of variation greater than 13%. A full description of methods used, as well as the atlas, the template that the atlas was derived from, and a masking file, can be found at Zenodo at https://doi.org/10.5281/zenodo.3555556. To our knowledge, this is the first MRI atlas created using Fischer 344 rats and will thus provide an appropriate neuroanatomical model for researchers working with this strain.

**HIGHLIGHTS:**

⍰ Open-access high-resolution anatomical MRI template for Fischer 344 rat brain.
⍰ Segmented atlas of 71 regions for use as a tool in Fischer 344 preclinical research paradigms.
⍰ Analysis of population variability of regional brain volumes.
⍰ Analysis of sex-differences in regional brain volumes
KEYWORDS: Fischer 344; Structural MRI; Segmentation; Rat brain template; Digital brain atlas; Sex-differences;

## 1. INTRODUCTION

In neuroscientific research involving preclinical rodent models, the ability to precisely identify and delineate anatomical brain regions is often a requirement.^1^ In the past, this identification was done using paper atlases such as Paxinos and Watson’s (P&W) *The Rat Brain in Stereotaxic Coordinates*^2^ However, with the increased prevalence of preclinical high-resolution magnetic resonance (MR) imaging in the past decade, digital neuroanatomical atlases have emerged in preclinical research as a tool for quick and accurate identification of anatomical regions in laboratory animals^3^. Digital atlases based on isotropic MRI provide flexibility over paper atlases by easily allowing interactive viewing of anatomical regions from arbitrary planes without distortion. This permits researchers to automatically record morphological information, such as the volume of brain regions, at the level of individual structures.^4,5^

A number of digital brain atlases for different rat strains have previously been published and disseminated,^6,7,8,9^ as shown in **Table 1**. The current study is motivated by the fact that there is currently no digital atlas for the Fischer 344 rat strain published in the literature or available online. Given that the Fischer 344 rat strain is commonly used in preclinical neuroscientific research,^10,11,12^ and given the emergence of novel transgenic rat models generated on a Fischer 344 background,^13,14,15^ a digital anatomical atlas of the Fischer 344 rat brain would be of value to the scientific community.

**Table 1.** Comparison between existing atlases and the current study.

| STUDY | Current Study | Papp et al. (2014) | Calabrese et al. (2013) | Schwarz et al. (2006) | Rumple et al. (2013) | Johnson et al. (2012) | Valdes-Hernandez et al. (2011) |
| --- | --- | --- | --- | --- | --- | --- | --- |
| RAT STRAIN | Fischer 344 | Sprague Dawley | Wistar | Sprague Dawley | Sprague Dawley | Wistar | Wistar |
| RESOLUTION OF T2-WEIGHTED MRI | 60 $\mu\text{m}^3$ | 39 $\mu\text{m}^3$ | 25 $\mu\text{m}^3$ | 190 $\mu\text{m}^3$ | 70 $\mu\text{m}^3$ for P5 and P14, 125 $\mu\text{m}^3$ for P72 | 25 $\mu\text{m}^3$ | 120x120x300 $\mu\text{m}$ |
| MODALITY | MR only | MR and DTI | MR and DTI | MR only | MR and DTI | MR, DTI, and histology | MR only |
| SAMPLE SIZE (n) | 41 mixed sex (24 male, 17 female) | 1 male | 5 males per timepoint | 97 males | 1 male and 1 female for P5 and P14, 5 females for P72 | 5 males | 5 males per timepoint |
| AVERAGE MR USED | Yes | No | Yes | Yes | Yes | Yes | Yes |
| ATLAS REGIONS | Whole Brain | Whole Brain | Whole Brain | Whole Brain | Whole Brain | Whole Brain | Whole Brain |
|  | In-vivo | Ex-vivo | Ex-vivo | In-vivo | In-vivo | Ex-vivo | In-vivo |
| LABEL METHOD | Stepwise manual segmentation based on T2 contrast on MR template, in conjunction with Paxinos and Watson atlas. | Semi-automatic (SNAP) and manual segmentation on MR. | MR average brains were aligned to conventional histological atlases. | Co-registration of average MR template to digital Paxinos and Watson histological atlas | Manual delineation in an iterative fashion based on T2 tissue contrast. Primarily delineated in coronal view, with sagittal and axial views used to verify segmentation. | Co-registration of average MR template to digital Paxinos and Watson histological atlas | Co-registration of average MR template to digital Paxinos and Watson histological atlas |
| NUMBER OF SEGMENTED STRUCTURES | 71 | 118 | 26 | 468 from co-registered Paxinos and Watson histological atlas | P5: 39 structures<br>P14: 45 structures<br>P72: 29 structures | 20 | 96 |
| **SUPPORTS**<br>**AUTOMATIC**<br>**DIGITAL**<br>**SEGMENTATION** | Yes | No | No | No | Yes | Yes: between<br>regions | No |
| **ANALYSIS OF**<br>**REGIONAL**<br>**VARIABILITY** | Yes: between<br>regions, and<br>between sex | No | Yes: between<br>regions | No | No | No | No |
| **FORMAT OF DIGITAL**<br>**ATLAS** | MINC and<br>NIfTI | NIfTI | NIfTI | Analyze<br>(AVW 7.5) | NRRD | Not specified | NIfTI |
| **PURPOSE OF STUDY** | Develop the<br>first tool for<br>automatic<br>digital<br>segmentation of<br>the Fischer 344<br>rat brain and to<br>examine<br>regional<br>variability of<br>Fischer rats,<br>allowing more<br>efficient<br>selection of<br>future<br>experiment<br>sample sizes. | Tool for spatial<br>analysis of<br>neuroanatomica<br>l location for<br>use in planning<br>and guidance of<br>experimental<br>procedures. | To establish a<br>timeline of<br>morphometric<br>changes and<br>variability<br>throughout<br>neurodevelopmen<br>t | Stereotaxic<br>MR template<br>with tissue<br>class<br>distribution<br>maps to<br>facilitate use<br>of fMRI<br>software in<br>tissue<br>segmentation<br>of brain data. | Digital atlas based<br>semi-automatic<br>segmentation to<br>increase<br>efficiency of<br>Sprague Dawley<br>analysis at P5,<br>P14, and P72. | Enhance<br>understanding<br>of<br>neuroanatom<br>y of the<br>Wistar rat and<br>offer a<br>collaborative<br>platform for<br>future rat<br>brain studies. | Template set<br>including<br>white and<br>grey matter<br>probabilistic<br>segmentation<br>for use in<br>fMRI<br>localization<br>in Wistar<br>rats. |

As seen in **Table 1**, there are several methods for generating whole brain atlases, as well as many different purposes for such atlases. The current atlas is the first reported atlas of the Fischer 344 rat brain. There are numerous advantages of the present atlas over currently existing rat brain atlases. Firstly, our atlas uses a relatively high number of scans (n=41), as well as using both males and female rats, to produce an average image which reduces bias from any single animal. In comparison, many other atlases use as few as 5 subjects or only male animals.^9,16,17^ Using a male-only generated atlas would be problematic when trying to apply the atlas to a mixed-sex study because there are clear sexual dimorphisms in specific brain regions, as documented thoroughly in the literature^18,19,20^ and in the current study. To our knowledge, the rat atlas by Schwarz et al. (2006) is the only one that uses more subjects (n=97 males) than in the current study.^21^ However, despite the higher number of rats used to generate their atlas, the image resolution was significantly lower than that of the atlas presented here.

All of the previously published rat atlases provide value to researchers working on different types of projects, whether they are interested in co-registering atlases to stereotaxic coordinates for use in surgery,^9^ to enhance our understanding of early neurodevelopment,^8^ or for semi-automatic segmentation of early-development rat brains.^15^ Aside from atlases generated for use in stereotaxic surgery which use a digitally co-registered P&W atlas, the present MRI atlas reports the highest number of delineated structures (n=71), which was possible due to the high isotropic resolution of the MR template image. Thus, the value of current study is that it reports a high-resolution, mixed-sex Fischer 344 rat atlas in MINC and NIfTI format for use in automatic structural segmentation and statistical analysis. Regional variation and structural sexual dimorphisms are also reported.

The practice of creating an MRI-based atlas generally involves generating a template brain using one of two approaches: a group-based approach or a single subject-based approach. The group-based approach takes a group of representative brains and creates an average image using deformation-based morphometric algorithms. In contrast, the single subject approach takes a single brain that is deemed to be most representative of the strain and uses that single brain as the template for the atlas. The benefit of the group-based approach is that it produces a high-resolution template while minimizing bias of a single subject, as well as capturing commonalities and variability in the population. Through statistical averaging, the average image can be up-sampled to a higher resolution than the initial subject images. A trade-off of the group-based approach is that small or highly variable regions can be lost due to statistical blurring. In contrast, the single subject approach, especially when used in conjunction with high-resolution ex-vivo imaging, can produce an exceptionally high-resolution MR image, but this is at the risk of including subject-specific anatomical abnormalities. In this work, we chose to develop an atlas of the Fischer 344 rat strain using a group-based approach. We also chose to use in-vivo MR scans in place of higher resolution ex-vivo scans, as the animals used are also part of an ongoing longitudinal study (unpublished). The described atlas contains 71 delineated structures throughout the whole brain, with an isotropic spatial resolution of 60 μm. The atlas was developed by manual segmentation of a group averaged rat brain and labels were assigned to each individual voxel to identify which voxels corresponded to which anatomical structure.

The resulting atlas can be utilized in conjunction with standard image processing pipelines used in cross-sectional and longitudinal neuroimaging studies^19,22^ and to provide structure-specific quantitative volumetric data at the single-subject level.^23^ By contrasting experimental groups against an appropriate control, this atlas would allow researchers to assess differences in regional brain volumes associated with an experimental or genetic manipulation. Furthermore, longitudinal assessment of volumetric changes is also possible by repeating the co-registration procedure using brain images obtained in the same sample at different timepoints. In practice, the above can be achieved by propagating this atlas onto experimental scans by co-registering individual rat brain scans to the template and using the Multiple Automatically Generated Template brain segmentation algorithm (MAGeT) described in Chakravarty et al, 2013.^22^ Importantly, one of the main strengths of this volumetric atlas is its compatibility with the Pydpiper and RMINC statistical software.^24,25^ The final atlas is provided in both MINC 2.0^26^ and NIfTI^27^ file formats to maximize utility for the community.

## 2. METHODS

### 2.1 Animal

Forty-four Fischer 344 rats were scanned at four months of age. Due to motion artefacts, 3 MRI scans were excluded from further analysis. The remaining forty-one adult Fischer 344 rats (24 males, 17 females) were free of major neuroanatomical abnormalities, based on visual inspection by a trained rater (DG) and were used to create the Fischer 344 atlas. At the time of scanning, the average age of the rats was 130 **±** 7 days, and the average weight was 282 **±** 60 g. The 41 rats were generated by two separate breeding colonies housed at the Douglas Hospital Research Centre’s Animal Facility in Montreal, Canada. Nine rats (3 F, 6 M) were the offspring of hemizygous male TgF344-AD (Tg) rats (acquired through a Material Transfer Agreement with the Terrence Town Laboratory at the University of Southern California) on a Fischer 344 background, bred with F344/NHsd wild-type (WT) females (010; Envigo Laboratories). Offspring from these breeders were a mixture of hemizygous Tg and homozygous wildtype (WT) rats, produced in approximately a 1:1 ratio per litter, of which only the WT Fischer 344 rats were used for the generation of this atlas. The remaining 32 rats (14 F, 16 M) were the products of F344/NHsd wild-type females bred with F344/NHsd wild-type males (010; Envigo Laboratories). All rats from both breeding schemes were genotyped using real time PCR to ensure none of the offspring for this study had incorporated the two transgenes from the TgF344-AD male breeders (Transnetyx, Cordova, TN). Since the wild-type rats used in this study came from two breeding schemes, an MRI-based structural analysis was conducted which showed no significant anatomical differences between the two schemes. Rats of the same sex were group housed (usually two per cage unless they showed signs of aggression), with ad libitum access to food and water. Animals were maintained under standard husbandry conditions on a 12/12 h light cycle, with lights on at 07:00 local time; the room temperature, relative humidity and air exchange were automatically controlled and monitored daily by animal facility staff.

#### Ethical approval

All applicable international, national, and/or institutional guidelines for the care and use of animals were followed. All animal procedures and experiments were approved by the McGill University Animal Care Committee (UACC) under protocol number 2016-7867.

### 2.2 MRI Acquisition

MRI data were acquired at the Douglas Centre d’Imagerie Cérébrale using a 7 Tesla Bruker Biospec 70/30 scanner (Bruker, Billerica MA, USA), with an 86 mm volumetric birdcage coil for transmission and a four-channel surface coil array for signal reception (Bruker). Rats were placed under anesthesia with a mixture of oxygen and isoflurane (4% iso during induction, then 2-4% for maintenance). The isoflurane level was adjusted to maintain a breathing rate between 45-65 breath/min throughout the procedure and warm air (37□) was blown into the bore of the scanner to maintain a constant body temperature. Animals were kept under anesthesia for 60 minutes while anatomical MRI, fMRI, and MRS data were acquired.

High-resolution 3D anatomical MR images were acquired using Rapid Acquisition with Relaxation Enhancement (RARE)^28^: TR = 325 ms, echo spacing = 10.8 ms, RARE factor = 6, effective echo time = 32.4 ms, FOV = 20.6 × 17.9 × 29.3 mm, matrix size = 256 × 180 x 157, slice thickness 17.9 mm (along the dorsal/ventral direction), readout along the rostral/caudal direction, scanner resolution = 114 μm isotropic, 19m35s acquisition time. Following the scan, animals were allowed to recover from anesthesia and returned to group housing.

### 2.3 Image Processing and Registration Pipelines

All images were processed in MINC format. Processing was performed using MINC-toolkit-v2^29^ and the Pydpiper module^25^ was used to co-register the processed images to produce a high-resolution co-registered average isotropic image.

Specimen registration was completed in an iterative process (**Figure 1)** to maximize the quality of the final template brain image using MINC-toolkit and the Pydpiper pipeline. First, raw anatomical MRI data from the scanner were exported in DICOM format and converted into MINC format for image pre-processing. Next, for each image an Otsu-threshold masking procedure was performed^30,^ followed by an N4 bias field correction^31^ in conjunction with the Otsu mask as weighting. The resulting intensity-normalized images were then run through the Pydpiper pipeline MBM.py which performs a co-registration of all input images to produce a group average image. The images initially underwent a rigid 6 parameter (LSQ6) alignment of rotations and translations to situate all images in a common space, followed by an affine 12 parameter (LSQ12) alignment which scales and shears the images pairwise to create an average. Finally, an iterative series of non-linear (nlin) alignments was performed to account for the remaining differences between brains. The output of this pipeline was upsampled to an isotropic resolution of 60 μm and manually masked to remove surrounding skull and non-brain tissue. Finally, the image was rotated 5 degrees about the z-axis to centre and square the image in the coronal view, resulting in an initial (first stage) group consensus average image.

**Figure 1:**
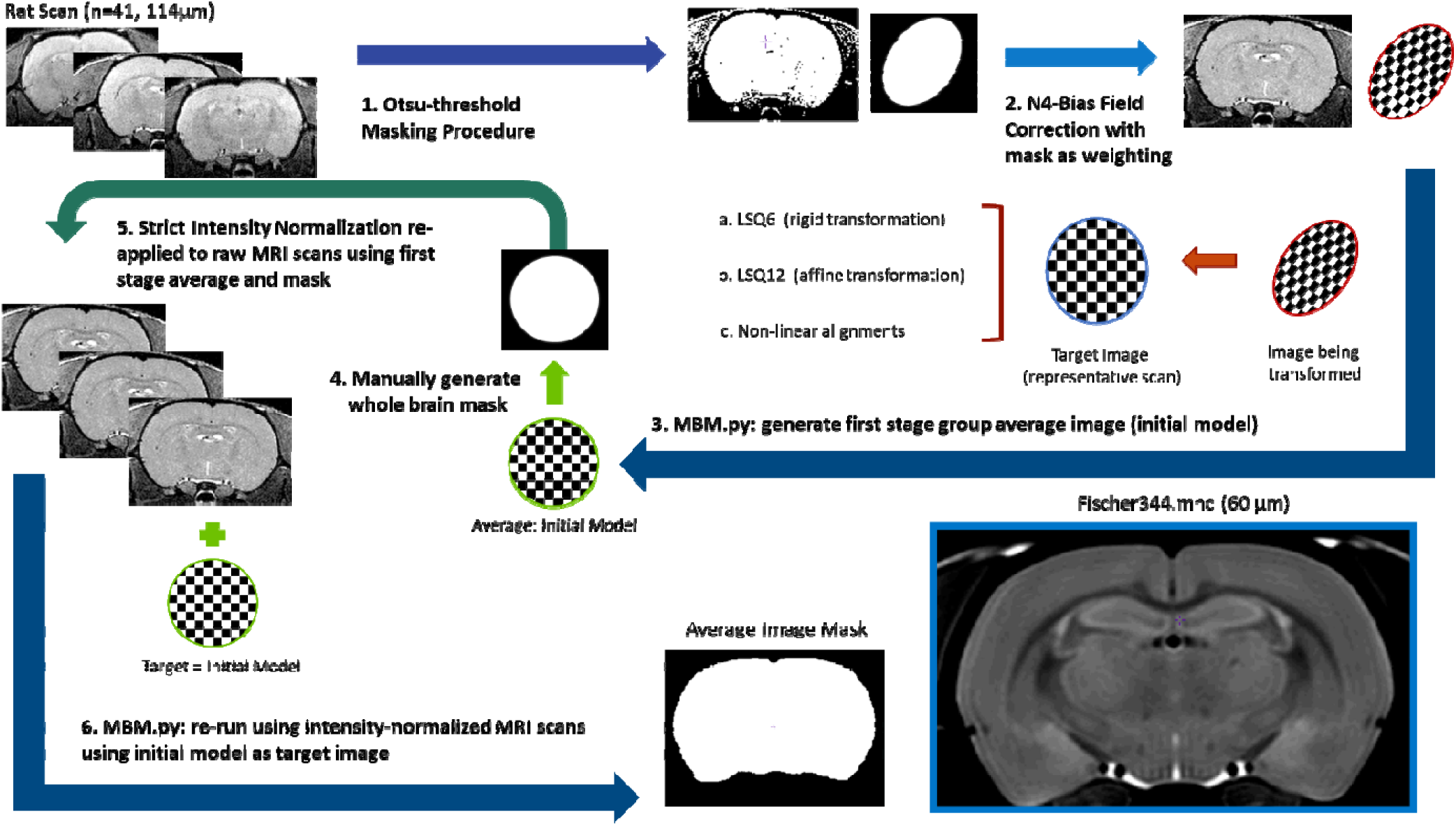
Flow chart demonstrating the iterative process used to produce the template image used for segmentation, as described in Section 2.3.

Using the image labelling tools within the minc-toolkit-v2 program Display, a whole-brain mask was manually generated based on this first stage group average image. Intensity normalizations were re-applied to the raw MRI image files by using the first stage average and whole-brain mask to strictly exclude non-brain regions from re-normalization. The resulting intensity normalized raw images were then run through the Pydpiper MBM.py pipeline using the first stage group average image as an initial model to increase image registration fidelity. The resulting average image with an isotropic resolution of 60 μm was then used as the final template image for the structural segmentation (**Figure 2)**.

**Figure 2:**
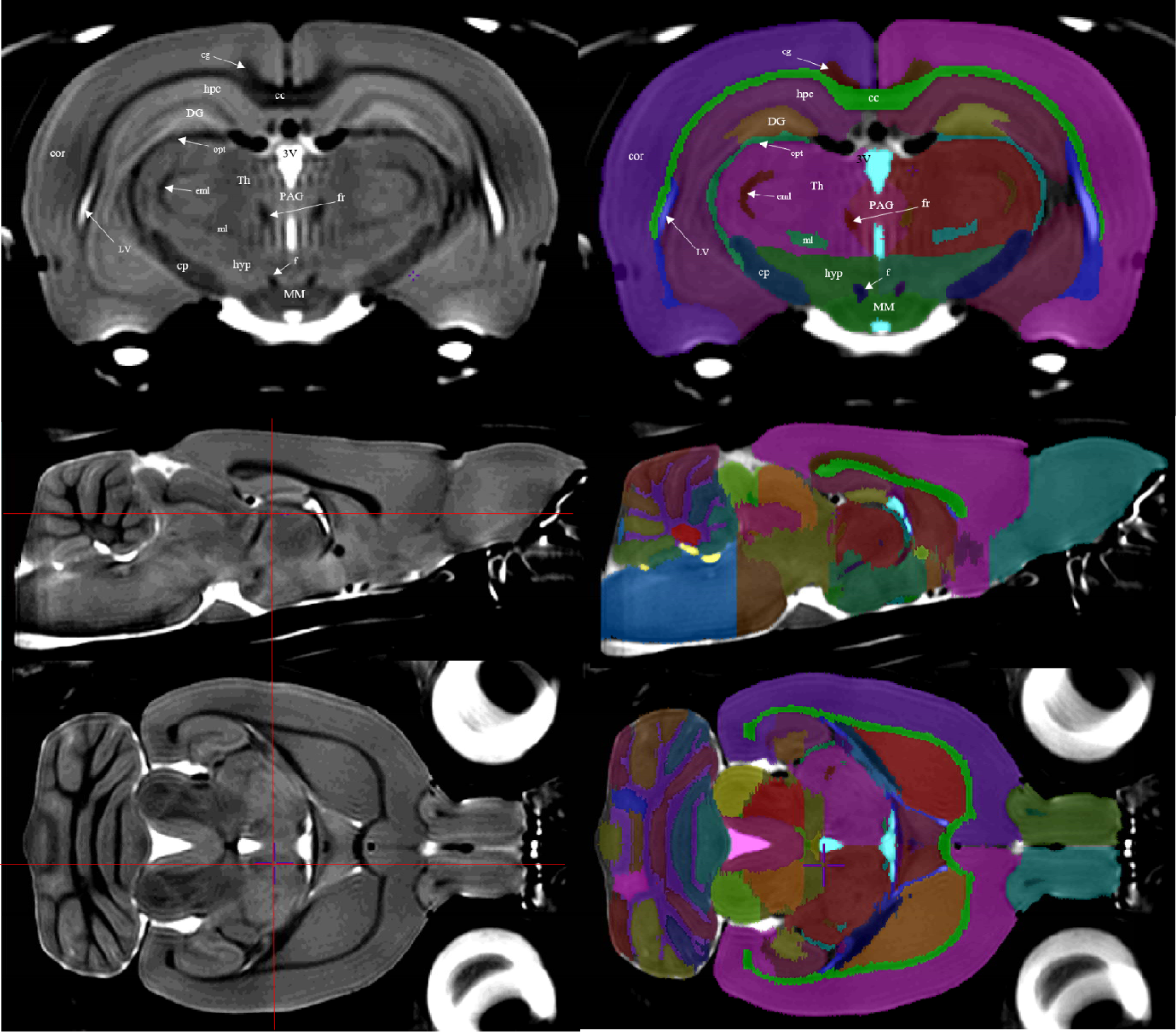
**Left column:** the averaged brain, which served as a template for structural delineation. Red crosshairs on transverse and sagittal view (middle and bottom) indicate the positions of the other planes. **Right column:** the atlas file is overlaid onto the template file. Delineation and refinement were primarily performed using the coronal sections, though further refinement was done in both the sagittal and axial planes. **Top row:** annotated over the left hemisphere of both coronal images are the structural regions that the label tags represent. **Middle** and **Bottom row** show transverse and sagittal views respectively; CC, corpus callosum and external capsule; cg, cingulum; cor, cortex; cp, cerebral peduncles; DG, dentate gyrus; eml, external medullary lamina; f, fornix; fr, fasciculus retroflexus; hyp, hypothalamus; hpc, hippocampal CA subfields; LV, lateral ventricle; ml, medial lemniscus; MM, mammillary bodies; opt, optic tract; PAG, periaqueductal gray; 3V, third ventricle.

### 2.4 Image Segmentation Protocol

Segmentation was performed by a single investigator (D.G.) based on the combined observations of T_2_-weighted tissue contrast as well as the P&W paper atlas^2^. Anatomical regions were delineated using the `Display` Software of the MINC-toolkit-v2 version 1.9.16 (https://github.com/BIC-MNI/minc-toolkit-v2).^29^ Positive grayscale contrast was used for all manual segmentation tasks. Regions were delineated on each coronal slice individually, and bilaterally, from the olfactory bulb to the first slice of spinal cord. In total, 71 anatomical structures or substructures were manually identified and included in the atlas. In certain specific regions, when boundaries were not obvious on basis of image contrast, the P&W atlas and anatomical landmarks were used to identify boundaries. Such regions are identified in further detail in the delineation section. The majority of the segmentation was done in the coronal view, however both the sagittal and axial views were used to ensure accuracy from other perspectives (**Figure 2**).

First, regions with significant and clear contrast from other regions such as the fibre tracts, ventricles, and caudoputamen were segmented. Next, peripheral brain regions such as the olfactory bulb, brainstem, and cerebellum were delineated. Inner brain nuclei that had poorer regional contrast were then delineated with the use of P&W Rat Brain atlas and by comparison with already delineated surrounding structures. A complete description of the methodology used to delineate individual structures can be found in the **Supplementary Methods Section**.

The resulting atlas MINC file contains numbered labels that correspond to each substructure, and is accompanied by a hierarchical Excel (.xlsx) file which lists the name and abbreviation of the structure associated with each label number, as well as its associated region (e.g. cerebellum, midbrain, hippocampus, etc.) and tissue type classification (GM, WM or CSF). This allows RMINC statistical software, for example, in a longitudinal deformation-based analysis, to identify regions of atrophy or growth by mapping the labeled atlas to the subject at various time points.

### 2.5 Statistical Analysis

The RMINC package in the R statistical environment was used to calculate the average volumes for each structure across all subjects (n=41) and to perform statistical comparisons between male (n=24) and female (n=17) volumes. The atlas label file was mapped to individual scans used in the analysis via the RMINC function `anatGetAll` which resulted in each voxel in each individual scan being uniquely labelled with a structural label. When mapping each rat brain to a common space, a MINC file containing the Jacobian determinant of each voxel required to scale the scan to the common space is generated. The RMINC function `anatGetAll` computes the structural volume for each region in mm^3^ by multiplying this scaling factor at each voxel by the voxel size and summing each value within each unique label. For the whole cohort average volume, the volume for each structure was averaged across the cohort, and standard deviation was computed. In addition to computing the mean and standard deviation of the volumes of each individual brain region across the cohort, we separately computed voxel-wise maps of the variability of the deformation fields in both male and female rats (see Supplementary Materials). Voxel-wise variability was expressed using the coefficient of variation of the relative Jacobians, and was calculated using RMINC,

For the sex-differences analysis, a two-sample, two-tailed Student’s T-test was performed with a Benjamini-Hochberg FDR correction (q=0.05) to control for multiple comparisons. The relative volume of each structure normalized to the volume of the subject’s total brain volume was computed to assay relative differences between male and female brain regions, reported in **Table 3**. In addition, the mean and standard deviations of regional volumes across the whole sample as well as for male and female rats are reported in **Table 2**.

**Table 2.** Regional volumes calculated for each label region as described in **Section 2.5 Statistical Analysis**, and averaged across brains. 71 delineated structures along with their corresponding mean and standard deviation (mm^**3**^) across all subjects (n=41), male rats (n=24) and female rats (n=17) are reported.

| VENTRICULAR<br>STRUCTURES | ALL SUBJECTS<br>MEAN + SD (mm<sup>3</sup>) | MALES<br>MEAN + SD (mm<sup>3</sup>) | FEMALES<br>MEAN + SD (mm<sup>3</sup>) |
| --- | --- | --- | --- |
| Aqueduct | $2.14 \pm 0.27$ | $2.29 \pm 0.23$ | $1.92 \pm 0.15$ |
| Fourth ventricle | $3.11 \pm 0.38$ | $3.37 \pm 0.28$ | $2.75 \pm 0.1$ |
| Lateral ventricle | $12.23 \pm 0.67$ | $12.66 \pm 0.51$ | $11.62 \pm 0.29$ |
| Third Ventricle | $5.17 \pm 0.26$ | $5.35 \pm 0.15$ | $4.92 \pm 0.17$ |
| **GREY MATTER**<br>**STRUCTURES** | **ALL SUBJECTS**<br>**MEAN + SD (mm<sup>3</sup>)** | **MALES**<br>**MEAN + SD (mm<sup>3</sup>)** | **FEMALES**<br>**MEAN + SD (mm<sup>3</sup>)** |
| Basal Forebrain | $15.84 \pm 0.89$ | $16.48 \pm 0.58$ | $14.95 \pm 0.28$ |
| Bed Nucleus of the Stria<br>Terminalis | $3.55 \pm 0.24$ | $3.74 \pm 0.09$ | $3.28 \pm 0.07$ |
| Caudoputamen | $75.72 \pm 3.26$ | $77.89 \pm 2.13$ | $72.66 \pm 1.73$ |
| Cerebellar Lobule 1/2 | $13.62 \pm 0.98$ | $14.35 \pm 0.47$ | $12.58 \pm 0.41$ |
| Cerebellar Lobule 10 | $3.31 \pm 0.23$ | $3.47 \pm 0.16$ | $3.1 \pm 0.12$ |
| Cerebellar Lobule 3 | $12.8 \pm 0.72$ | $13.27 \pm 0.52$ | $12.12 \pm 0.34$ |
| Cerebellar Lobule 4/5 | $33.11 \pm 1.94$ | $34.42 \pm 1.32$ | $31.25 \pm 0.79$ |
| Cerebellar Lobule 6 | $11.9 \pm 0.92$ | $12.51 \pm 0.62$ | $11.04 \pm 0.47$ |
| Cerebellar Lobule 7 | $4.49 \pm 0.28$ | $4.67 \pm 0.2$ | $4.24 \pm 0.17$ |
| Cerebellar Lobule 8 | $4.88 \pm 0.36$ | $5.1 \pm 0.28$ | $4.57 \pm 0.22$ |
| Cerebellar Lobule 9 | $10.65 \pm 0.76$ | $11.21 \pm 0.4$ | $9.85 \pm 0.25$ |
| Cochlear Nucleus | $1.91 \pm 0.11$ | $2 \pm 0.05$ | $1.78 \pm 0.04$ |
| Copula | $3.63 \pm 0.28$ | $3.85 \pm 0.12$ | $3.32 \pm 0.07$ |
| Cortex | $704.44 \pm 38.59$ | $730.82 \pm 25.98$ | $667.21 \pm 15.35$ |
| Crus 1 Ansiform Lobule | $24.63 \pm 1.44$ | $25.6 \pm 0.98$ | $23.25 \pm 0.58$ |
| Crus 2 Ansiform Lobule | $10.69 \pm 0.67$ | $11.11 \pm 0.54$ | $10.1 \pm 0.29$ |
| Dentate Gyrus | $23.14 \pm 1.58$ | $24.29 \pm 0.9$ | $21.53 \pm 0.61$ |
| Dentate Nucleus | $2 \pm 0.12$ | $2.09 \pm 0.08$ | $1.88 \pm 0.03$ |
| Entopeduncular Nucleus | $0.75 \pm 0.04$ | $0.78 \pm 0.03$ | $0.71 \pm 0.02$ |
| Fastigial Nucleus | $2.35 \pm 0.13$ | $2.45 \pm 0.07$ | $2.21 \pm 0.05$ |
| Flocculus | $9.99 \pm 0.73$ | $10.45 \pm 0.48$ | $9.35 \pm 0.49$ |
| Globus Pallidus | $7.91 \pm 0.39$ | $8.16 \pm 0.28$ | $7.56 \pm 0.23$ |
| Hindbrain | $165.67 \pm 10.55$ | $173.39 \pm 5.91$ | $154.77 \pm 3.57$ |
| Hippocampal Formation | $67.77 \pm 3.86$ | $70.48 \pm 1.97$ | $63.94 \pm 2.27$ |
| Hypothalamus | $44.08 \pm 2.54$ | $45.94 \pm 1.46$ | $41.44 \pm 0.69$ |
| Inferior Colliculus | $25.39 \pm 1.21$ | $26.16 \pm 0.9$ | $24.3 \pm 0.61$ |
| Interposed Nucleus | $2.59 \pm 0.13$ | $2.69 \pm 0.07$ | $2.45 \pm 0.04$ |
| Lateral Septum | $13.04 \pm 0.65$ | $13.49 \pm 0.41$ | $12.4 \pm 0.28$ |
| Mammillary bodies | $2.1 \pm 0.12$ | $2.18 \pm 0.09$ | $2 \pm 0.07$ |
| Medial Septum | $2.08 \pm 0.11$ | $2.15 \pm 0.07$ | $1.97 \pm 0.05$ |
| Median Preoptic Nucleus | $0.08 \pm 0.01$ | $0.08 \pm 0.01$ | $0.07 \pm 0$ |
| Midbrain | $64.77 \pm 4.08$ | $67.76 \pm 2.41$ | $60.55 \pm 0.97$ |
| Nucleus Accumbens | $13.34 \pm 0.56$ | $13.69 \pm 0.41$ | $12.84 \pm 0.32$ |
| Olfactory Nuclei | $131.13 \pm 8.38$ | $137.18 \pm 4.89$ | $122.59 \pm 2.96$ |
| Paraflocculus | $14.93 \pm 1$ | $15.55 \pm 0.66$ | $14.06 \pm 0.69$ |
| Paramedian Lobule | $10.97 \pm 0.7$ | $11.48 \pm 0.43$ | $10.26 \pm 0.21$ |
| Periaqueductal Grey | $15.26 \pm 0.63$ | $15.7 \pm 0.4$ | $14.65 \pm 0.27$ |
| Pons | $45.38 \pm 2.89$ | $47.31 \pm 2.13$ | $42.66 \pm 0.97$ |
| Simple Lobule | $22.98 \pm 1.26$ | $23.85 \pm 0.68$ | $21.74 \pm 0.71$ |
| Subfornical Organ | 0.13 ± 0.01 | 0.14 ± 0.01 | 0.12 ± 0.01 |
| Substantia Nigra | 4.71 ± 0.29 | 4.91 ± 0.22 | 4.44 ± 0.09 |
| Superior Colliculus | 27.33 ± 1.55 | 28.39 ± 0.96 | 25.83 ± 0.77 |
| Thalamus | 74.18 ± 3.55 | 76.59 ± 2.22 | 70.78 ± 1.84 |
| Ventral Pallidum | 4.17 ± 0.2 | 4.31 ± 0.12 | 3.97 ± 0.08 |
| **WHITE MATTER**<br>**STRUCTURES** | **ALL SUBJECTS**<br>**MEAN + SD (mm<sup>3</sup>)** | **MALES**<br>**MEAN + SD (mm<sup>3</sup>)** | **FEMALES**<br>**MEAN + SD (mm<sup>3</sup>)** |
| Anterior Part of the<br>Anterior Commissure | 2.69 ± 0.13 | 2.79 ± 0.08 | 2.56 ± 0.06 |
| Cerebellar White Matter<br>and Arbor Vitae | 47.55 ± 2.77 | 49.59 ± 1.51 | 44.67 ± 0.96 |
| Cerebral Peduncle | 6.29 ± 0.34 | 6.51 ± 0.27 | 5.97 ± 0.12 |
| Cingulum | 7.12 ± 0.44 | 7.43 ± 0.27 | 6.69 ± 0.2 |
| Commissure of the Inferior<br>Colliculus | 0.63 ± 0.02 | 0.64 ± 0.02 | 0.61 ± 0.02 |
| Commissure of the<br>Superior Colliculus | 0.32 ± 0.02 | 0.33 ± 0.02 | 0.3 ± 0.01 |
| Corpus Callosum and<br>External Capsule | 64.4 ± 3.66 | 66.9 ± 2.18 | 60.89 ± 2.08 |
| External Medullary Lamina | 1.55 ± 0.09 | 1.62 ± 0.05 | 1.46 ± 0.03 |
| Fasciculus Retroflexus | 0.36 ± 0.01 | 0.37 ± 0.01 | 0.34 ± 0.01 |
| Fimbria | 7.05 ± 0.45 | 7.36 ± 0.24 | 6.6 ± 0.24 |
| Fornix | 0.9 ± 0.04 | 0.93 ± 0.02 | 0.85 ± 0.01 |
| Internal Capsule | 15.59 ± 0.88 | 16.17 ± 0.61 | 14.77 ± 0.4 |
| Intrabulbar Part of Anterior | 1.66 ± 0.08 | 1.71 ± 0.05 | 1.59 ± 0.05 |
| Commissure |  |  |  |
| Lateral Olfactory Tract | $1.08 \pm 0.05$ | $1.12 \pm 0.03$ | $1.03 \pm 0.03$ |
| Mammillothalamic Tract | $0.46 \pm 0.02$ | $0.47 \pm 0.01$ | $0.44 \pm 0.01$ |
| Optic Chiasm | $0.52 \pm 0.06$ | $0.57 \pm 0.04$ | $0.46 \pm 0.02$ |
| Optic Tract | $6.19 \pm 0.44$ | $6.52 \pm 0.23$ | $5.73 \pm 0.16$ |
| Posterior Commissure | $0.28 \pm 0.02$ | $0.3 \pm 0.01$ | $0.27 \pm 0.01$ |
| Posterior Part of the<br>Anterior Commissure | $0.66 \pm 0.03$ | $0.69 \pm 0.02$ | $0.62 \pm 0.01$ |
| Stria Medullaris of the<br>Thalamus | $0.67 \pm 0.03$ | $0.69 \pm 0.02$ | $0.64 \pm 0.01$ |
| Stria Terminalis | $2.11 \pm 0.14$ | $2.21 \pm 0.09$ | $1.98 \pm 0.06$ |
| Superior Thalamic<br>Radiation | $0.89 \pm 0.05$ | $0.92 \pm 0.03$ | $0.85 \pm 0.03$ |

**Table 3.**
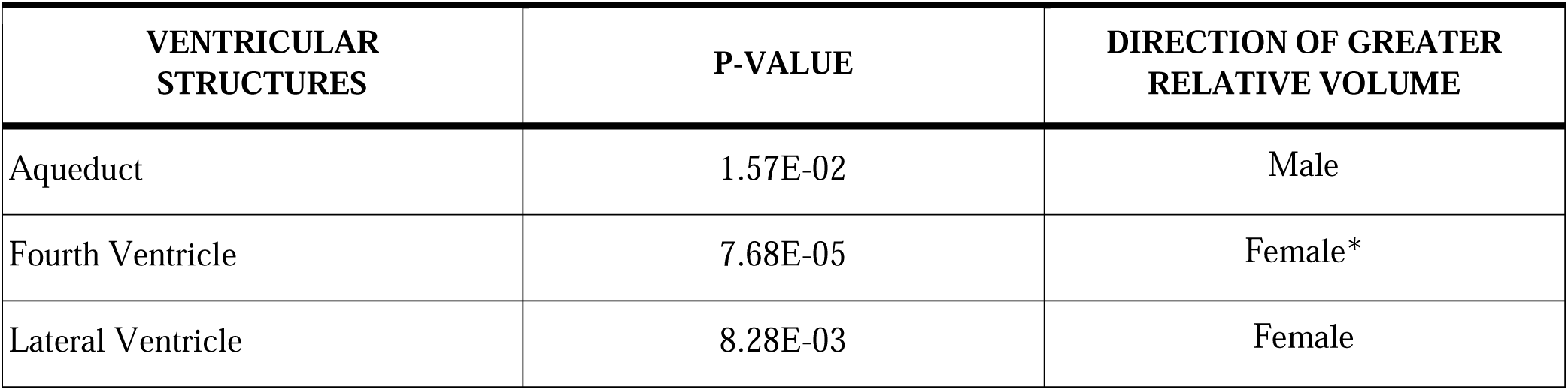
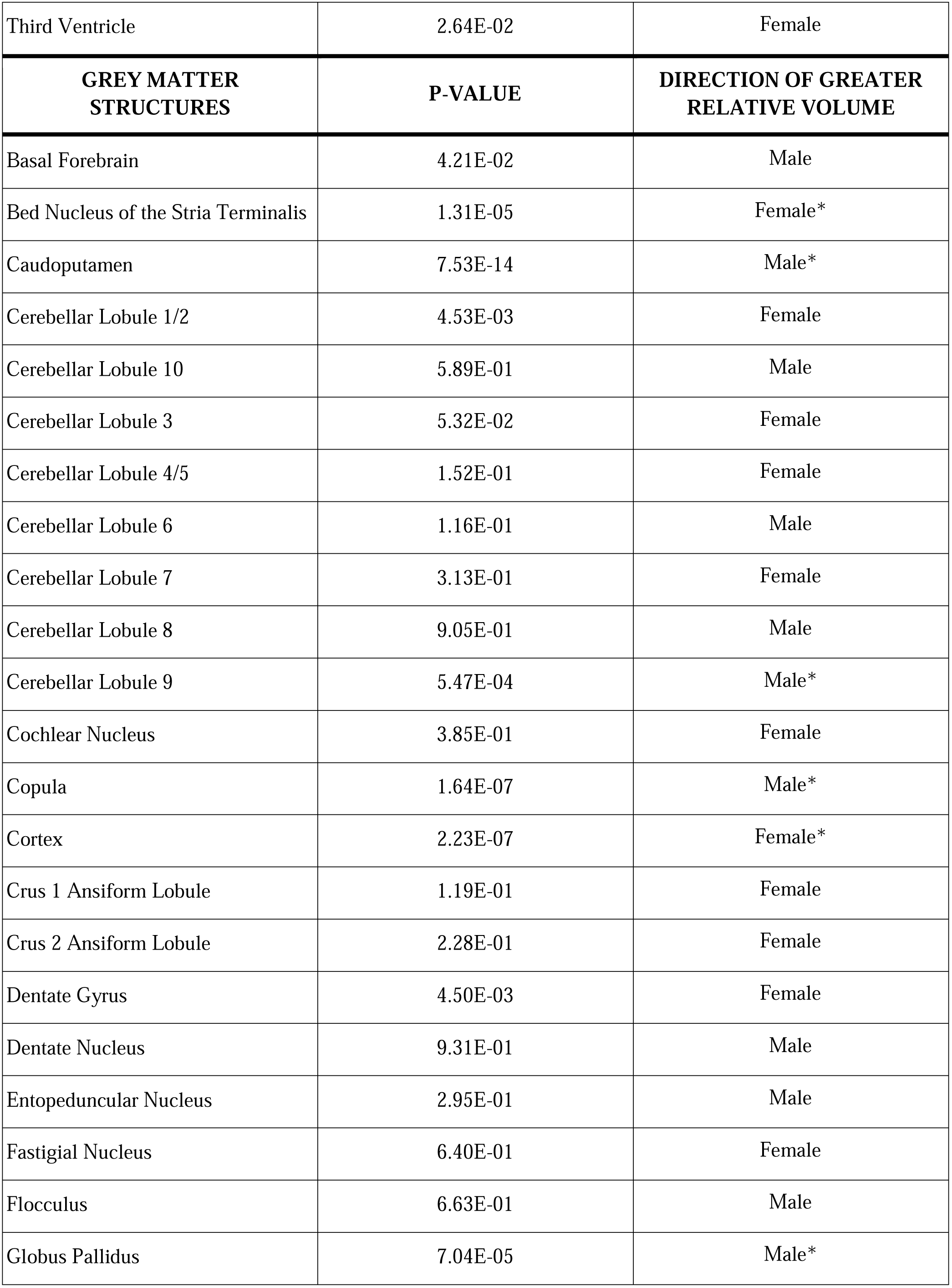
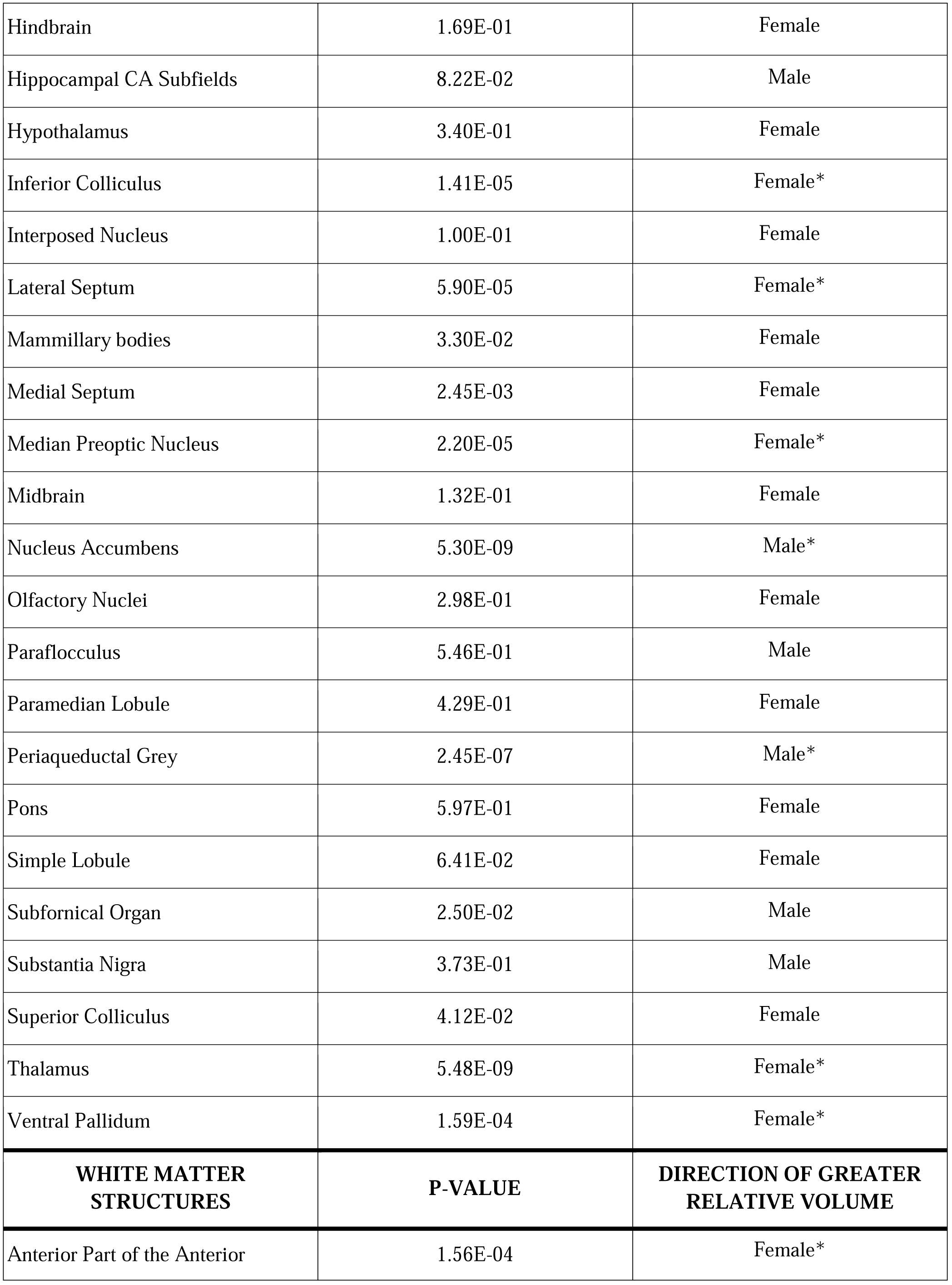
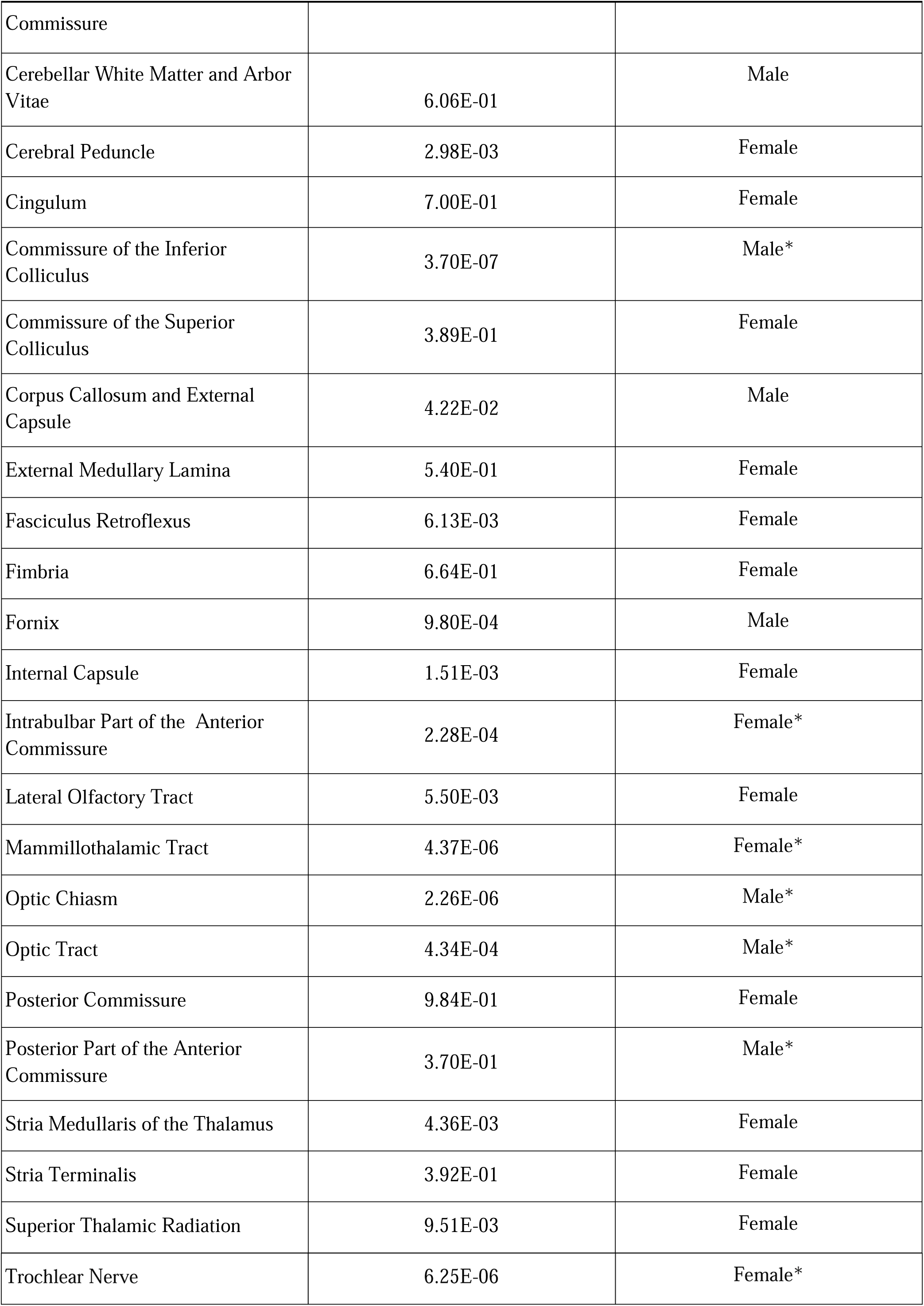
Two sample, two tailed t-test between the relative volumes of male (n=24) and female (n=17) groups. The second column indicates the p-value obtained from the t-test. Significance between relative size of regions between males and females is denoted by an asterisk in the third column. The third column indicates the direction of greater relative volume. Of specific interest is the cortex with females having significantly (p<0.0001) larger relative cortex than males.

| VENTRICULAR STRUCTURES | P-VALUE | DIRECTION OF GREATER RELATIVE VOLUME |
| --- | --- | --- |
| Aqueduct | 1.57E-02 | Male |
| Fourth Ventricle | 7.68E-05 | Female* |
| Lateral Ventricle | 8.28E-03 | Female |

**Table 3.** Two sample, two tailed t-test between the relative volumes of male (n=24) and female (n=17) groups.
| Third Ventricle | 2.64E-02 | Female |
| --- | --- | --- |
| **GREY MATTER**<br>**STRUCTURES** | **P-VALUE** | **DIRECTION OF GREATER**<br>**RELATIVE VOLUME** |
| Basal Forebrain | 4.21E-02 | Male |
| Bed Nucleus of the Stria Terminalis | 1.31E-05 | Female* |
| Caudoputamen | 7.53E-14 | Male* |
| Cerebellar Lobule 1/2 | 4.53E-03 | Female |
| Cerebellar Lobule 10 | 5.89E-01 | Male |
| Cerebellar Lobule 3 | 5.32E-02 | Female |
| Cerebellar Lobule 4/5 | 1.52E-01 | Female |
| Cerebellar Lobule 6 | 1.16E-01 | Male |
| Cerebellar Lobule 7 | 3.13E-01 | Female |
| Cerebellar Lobule 8 | 9.05E-01 | Male |
| Cerebellar Lobule 9 | 5.47E-04 | Male* |
| Cochlear Nucleus | 3.85E-01 | Female |
| Copula | 1.64E-07 | Male* |
| Cortex | 2.23E-07 | Female* |
| Crus 1 Ansiform Lobule | 1.19E-01 | Female |
| Crus 2 Ansiform Lobule | 2.28E-01 | Female |
| Dentate Gyrus | 4.50E-03 | Female |
| Dentate Nucleus | 9.31E-01 | Male |
| Entopeduncular Nucleus | 2.95E-01 | Male |
| Fastigial Nucleus | 6.40E-01 | Female |
| Flocculus | 6.63E-01 | Male |
| Globus Pallidus | 7.04E-05 | Male* |
| Hindbrain | 1.69E-01 | Female |
| Hippocampal CA Subfields | 8.22E-02 | Male |
| Hypothalamus | 3.40E-01 | Female |
| Inferior Colliculus | 1.41E-05 | Female* |
| Interposed Nucleus | 1.00E-01 | Female |
| Lateral Septum | 5.90E-05 | Female* |
| Mammillary bodies | 3.30E-02 | Female |
| Medial Septum | 2.45E-03 | Female |
| Median Preoptic Nucleus | 2.20E-05 | Female* |
| Midbrain | 1.32E-01 | Female |
| Nucleus Accumbens | 5.30E-09 | Male* |
| Olfactory Nuclei | 2.98E-01 | Female |
| Paraflocculus | 5.46E-01 | Male |
| Paramedian Lobule | 4.29E-01 | Female |
| Periaqueductal Grey | 2.45E-07 | Male* |
| Pons | 5.97E-01 | Female |
| Simple Lobule | 6.41E-02 | Female |
| Subfornical Organ | 2.50E-02 | Male |
| Substantia Nigra | 3.73E-01 | Male |
| Superior Colliculus | 4.12E-02 | Female |
| Thalamus | 5.48E-09 | Female* |
| Ventral Pallidum | 1.59E-04 | Female* |
| **WHITE MATTER**<br>**STRUCTURES** | **P-VALUE** | **DIRECTION OF GREATER**<br>**RELATIVE VOLUME** |
| Anterior Part of the Anterior | 1.56E-04 | Female* |
| Commissure |  |  |
| Cerebellar White Matter and Arbor Vitae | 6.06E-01 | Male |
| Cerebral Peduncle | 2.98E-03 | Female |
| Cingulum | 7.00E-01 | Female |
| Commissure of the Inferior Colliculus | 3.70E-07 | Male* |
| Commissure of the Superior Colliculus | 3.89E-01 | Female |
| Corpus Callosum and External Capsule | 4.22E-02 | Male |
| External Medullary Lamina | 5.40E-01 | Female |
| Fasciculus Retroflexus | 6.13E-03 | Female |
| Fimbria | 6.64E-01 | Female |
| Fornix | 9.80E-04 | Male |
| Internal Capsule | 1.51E-03 | Female |
| Intrabulbar Part of the Anterior Commissure | 2.28E-04 | Female* |
| Lateral Olfactory Tract | 5.50E-03 | Female |
| Mammillothalamic Tract | 4.37E-06 | Female* |
| Optic Chiasm | 2.26E-06 | Male* |
| Optic Tract | 4.34E-04 | Male* |
| Posterior Commissure | 9.84E-01 | Female |
| Posterior Part of the Anterior Commissure | 3.70E-01 | Male* |
| Stria Medullaris of the Thalamus | 4.36E-03 | Female |
| Stria Terminalis | 3.92E-01 | Female |
| Superior Thalamic Radiation | 9.51E-03 | Female |
| Trochlear Nerve | 6.25E-06 | Female* |

### 2.6 Tissue Probability Maps

Following the generation of the atlas, a manually derived tissue classification label image was generated by assigning a classification value of 1, 2, 3 to all structures classified in the hierarchy file as GM, WM, and CSF, respectively. Any remaining pixels inside the brain mask that were not assigned to one of the segmented structures were assigned as either CSF (classification value 3, if its pixel intensity was greater than 2500), or other (classification value 4, if its pixel intensity was 2500 or less). The resulting GM/WM/CSF/other classification image was used as an initial prior for the generation of a tissue probability map using Atropos.^32^ The primary input to the Atropos function was the average brain template, and the manually segmented GM/WM/CSF/other prior classification image was given a weighting of 0.5 in the optimization.

### 2.7 Data Sharing

The final anatomical segmentation volume along with the template brain, corresponding label descriptions, and a brain masking volume were each converted from MINC format into NIfTI format using the minc-toolkit `mnc2nii` function. All of the above are available for download and free use through Zenodo at https://doi.org/10.5281/zenodo.3555556 in both NIfTI and the MINC 2.0 format – compatible with all MINC tools, as well as Pydpiper and RMINC software.

## 3. RESULTS

In this Fischer 344 rat atlas, we delineated 71 anatomical structures comprising the ventricular system, grey matter, and white matter tracts. The level of detail in the averaged brain template is exemplified along with an overview of the anatomical delineations in **Figures 2** and **3**. It should be noted that good image contrast in the template has allowed for relatively detailed delineations in the cerebellum (**Figure. 4**). Based on the anatomical labels, and using the methods described in **Section 2.5**, regional volumes for regions distinguishable based on T2-weighted contrast were calculated for each of the 41 brains. Average volumes for each region, as well as the level of variability within and across male and female animals are summarized in **Table 2**.

**Figure 3:**
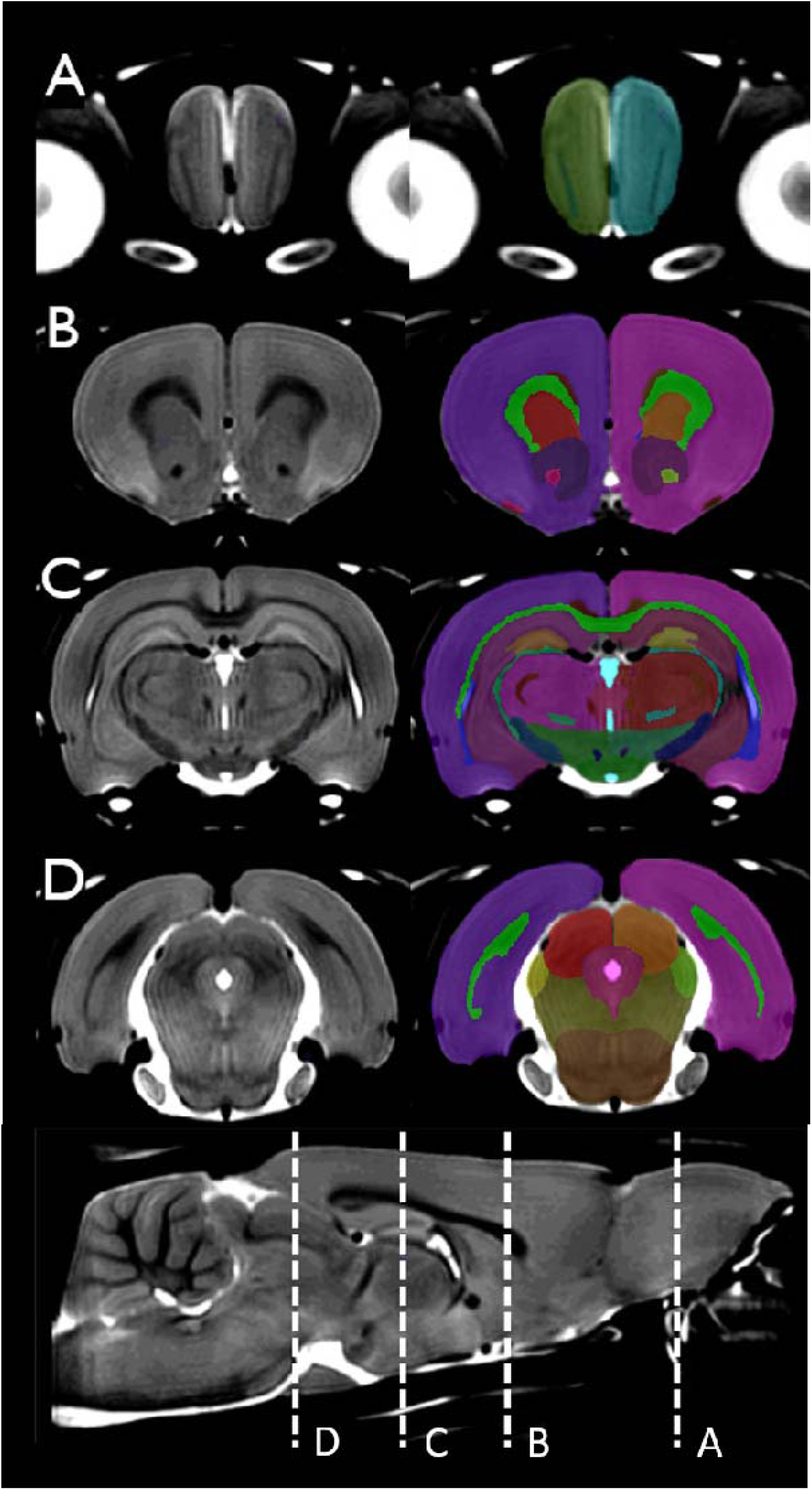
In the **left column** is the template brain, and in the **right column** is the template brain with the atlas file superimposed. This figure shows a series of coronal slices at different regions of the neocortex. The positions of representative slices are marked with dashed lines in a mid-sagittal slice with labels A-D.

**Figure 4:**
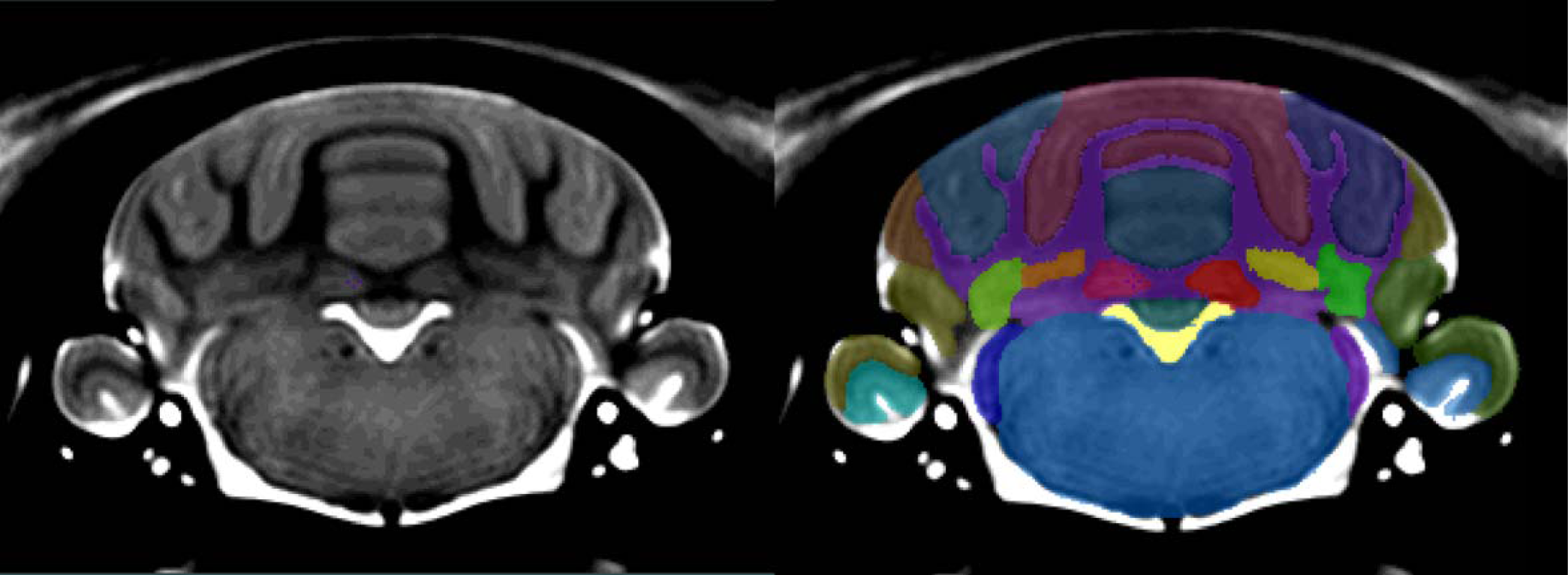
On the **left** is the template brain, and on the **right** is the template brain with the atlas file superimposed. Clear boundaries are identifiable between all of the major lobes. White matter and Arbor Vitae were indistinguishable from each other based on the resolution of the image. All the hindbrain nuclei were delineated as one structure due to poor resolution and low contrast levels in the hindbrain.

On average, the variability of brain regions across all samples expressed in terms of percent of the region volume was 5% (**Figure 5**), indicating that there are fairly small regional variations across four-month-old Fischer 344 rats. In the ventricular system the variability reached 12.2% (4^th^ ventricle), which can possibly be attributed to differences in hydration levels between subjects resulting in variant CSF levels.

**Figure 5:**
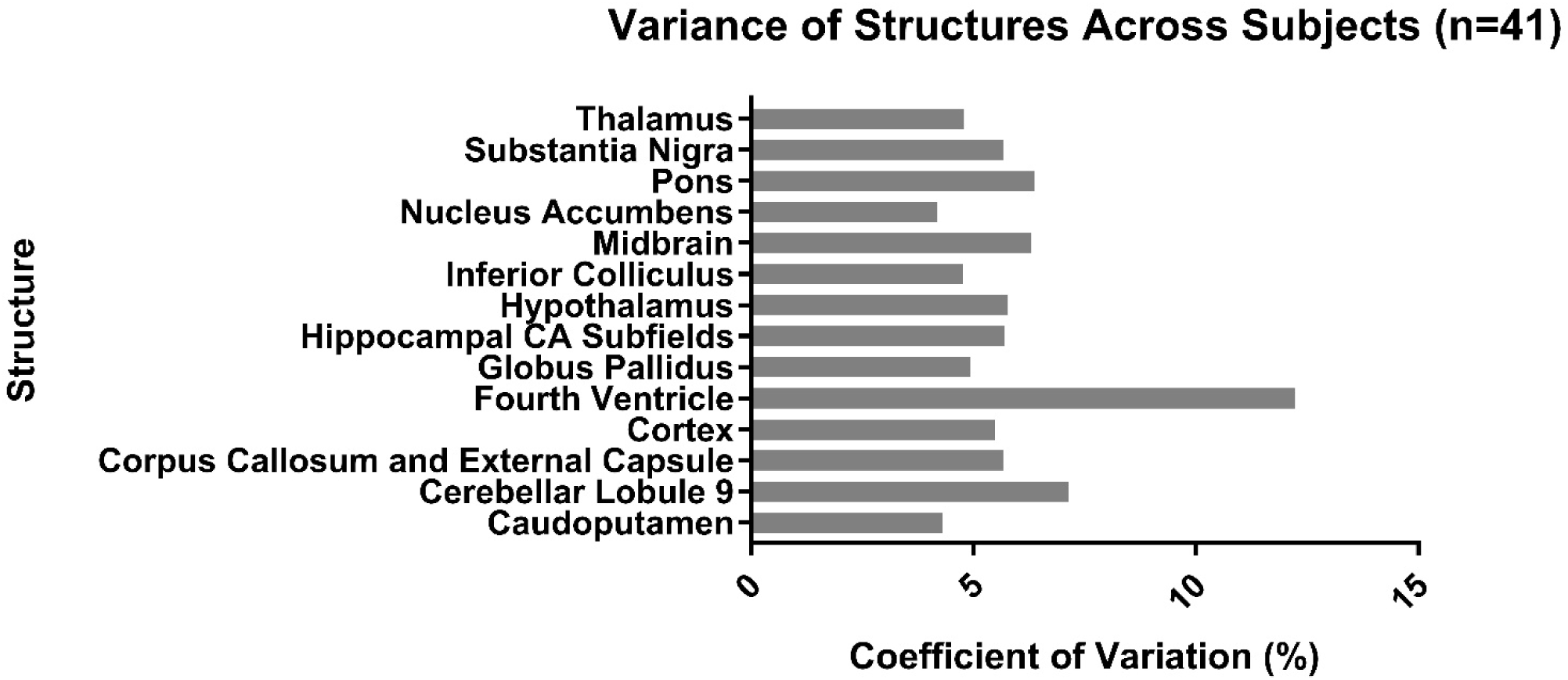
Comparison of the coefficient of variation (CV) for selected structures across all 41 subjects. The volume of most of the 71 structures varies between 4% and 8% across subjects. As expected, the ventricular system displays increased variation across subjects (Fourth Ventricle: 12.2%).

We performed two different analyses to compare structural volumes^23^ between male and female rats. First, we assessed the absolute volumes of each region to evaluate which regions were significantly different in size. Next, we looked at relative volumes (absolute regional volume divided by whole brain volume) to evaluate differences between the relative regional volumes between sexes. When evaluating absolute size differences between sexes, all regions were significantly larger in males relative to females (see **Supplemental Table 1** for more details). This is understandable due to animal size differences—the mean total volume of the male brains was 2002 ± 64 mm^3^ while the mean total volume of female brains was 1817 ± 36 mm^3^. In contrast, when evaluating relative volumes, certain volumes were significantly larger in females. **Table 3** reports the results of a two-tailed t-test with correction for multiple comparisons between the relative regional volumes between male and female Fischer 344 rats. One specific finding of interest is that females have a relatively larger neocortex (p<0.001).

The voxel-wise maps of the coefficient of variation of the deformation fields are shown in the Supplementary Materials for both male (Figure S3) and female (Figure S4) rats. Based on these figures, brain regions associated with the largest inter-individual variability in can be identified. Nowhere in the brain does the coefficient of variation of the deformation fields exceed +/-2%.

The tissue probability maps generated using Atropos are shown in the Supplementary Materials (Figure S5), and clearly illustrate the spatial distributions of GM, WM, CSF and other tissue.

## 4. DISCUSSION

In the present study we developed a 60 μm isotropic template image of the normative Fischer 344 rat brain, from which we created an atlas comprised of 71 structures and provided a detailed basis for delineation for each structure. We provide these tools, along with a template brain masking file in both the MINC 2.0 and NIfTI imaging file format, as open-access tools available to researchers to validate as well as to facilitate highly detailed structural analyses.

This paper presents a tool for preclinical researchers to identify brain structures within MR images of the Fischer 344 rat brain in a digital format, enabling rapid semi-automated structural analyses to be performed. Additionally, due to the automated nature of the identification of brain regions using software such as MAGeT or RMINC, segmentation bias is reduced across subjects compared with manual individual segmentation. As rat brain development is completed by the end of adolescence, or post-natal day 45, this tool can be reliably used on most adult Fischer 344 MR images. In order to minimize segmentation inaccuracies, when mapping a set of Fischer 344 volumes to our template image, volumes should be within 10% of the template image volume: 1925.59 ± 106.80 mm^3^.^16^

Though other atlases such as the gold standard Paxinos and Watson histological atlas may be able to provide more precise identification of smaller nuclei that is not attainable on an in-vivo MRI based atlas such as that presented here, this work provides the rodent neuroimaging community with the first high-field (7T) *in vivo* Fischer 344 rat brain MRI atlas, a high resolution template isotropic image, and label hierarchy that can be tailored to the desired anatomical specificity of analysis. Of the few *in vivo* MRI atlases that presently exist,^7,34,35,36^ ours is the first to use both male and female rats and a comparatively large sample size (this study, n-41; next largest is n=30, smallest is n=5). The result is an MRI atlas with segmentations generated from a relatively diverse cohort, which increases its applicability to studies examining both sexes.

In addition to the development of a tool for morphological analysis, we report an analysis of the variability of regional brain. One particular finding of interest in the sex-difference analysis was that female Fischer 344 rats have relatively larger neocortices **(Table 3)**. Many of the other regions that were also found to be significantly larger in females were fibre tracts involved in cortico-cortical communication such as the internal capsule, anterior commissure and the commissures of the colliculi. These findings are logical: since the neocortex is relatively larger, the fibre tracts that connect the different cortical regions would also be expected to be larger.

Furthermore, the absolute volumes of each anatomical region across the whole group, as well as between sexes reported in **Table 2** provide valuable information to researchers attempting to plan future studies. This data will allow researchers to better plan sample sizes when performing a longitudinal study of regional atrophy, for example. Highly detailed data on the structural volumes of Fischer 344 rat brains is not documented in the literature, and thus this analysis provides novel value for future investigations.

This resource was developed using scans taken from 41 four-month-old male and female Fischer 344 wild-type rats weighing 282g ± 60g, and therefore provides structural information relevant to mature adult rats. 32 of the subjects were wild-type (WT/WT) offspring from homozygous wild-type breeders, and 9 were wild-type (WT/WT) offspring from a hemizygous (WT/Tg) transgenic male bred with a homozygous wild-type female (WT/WT). A structural comparison using the Pydpiper pipeline confirmed that there is no structural variance between the wild-type rats that came from different lineages.

It is also important to note that the rats used to generate the atlas were bred in-house using breeders purchased from Envigo and/or the Terrence Town Laboratory rather than acquiring them directly from a vendor, due to these rats being part of a larger longitudinal study. While the in-house breeding may represent a minor limitation regarding reproducibility, it should not play any larger a role than the other factors known to introduce variation in animal research; housing conditions, handling, shipping conditions, litter size, and maternal care, among others, have been shown to affect genetic and phenotypic presentation in rodents, not to mention the thoroughly documented physiological differences within the same rat strain obtained from different vendors (i.e. Sprague-Dawley obtained from Charles River versus Envigo).^37,38,39,40,41^ While no studies exist that directly compare neuroanatomical differences between rats subjected to any of the above confounding variables, care should be taken when choosing the rat strain, vendor, housing conditions, etc., when attempting to use resources generated from a specific population having experienced a facility-specific environment.

Finally, regarding the generalizability of this Fischer 344 atlas to other rat strains, many studies have successfully applied atlases generated from strains differing from their experimental rats, likely in part due to the lack of an appropriate MRI atlas for their own rat strain and/or research question.^16,42,43,44^ In the literature, the general consensus appears to be that any particular atlas, including that designed by Paxinos and Watson, may be applied to both sexes, different strains, etc., provided there are no gross anatomical abnormalities, the age and weight range of the subjects is comparable, and volumes should be within 10% of the template image volume.^2,16,35,42^

That said, volume differences in both whole brain and specific structures have been shown between rat strains.^42^ For example, Welniak-Kaminska et al. found that whole brain volumes of Brown Norway (BN), Wistar (WI), and Warsaw Wild Captive Pisula Strykek (WWCPS) rats were 1832±25.9 mm^3^, 1971±22.7 mm^3^, and 1712±28.5 mm^3^, respectively, and that their body weight (strongly related to brain volume) also differed significantly, despite all rats being comparable ages (56 to 63 days). The same study showed significant differences in entorhinal cortex, amygdala, and hippocampal volumes, among others. Even within-strain differences can be seen by comparing two studies citing whole brain volumes of Wistar rats: although the MRI template set provided by Valdéz-Hernández et al.^7^ was created using Wistar rats of a similar age and weight range to those studied by Welniak-Kaminska et al., Valdéz-Hernández et al. cites a whole brain volume of 1764.92 ± 85.57 mm^3^, compared to 1971±22.7 mm^3^ quoted by Welniak-Kaminska, which is outside the 10% difference in volume guidelines stated previously. For comparison, the whole brain volume of Fischer rats in this study is 1925±106.80 mm^3^, which is within 10% of the whole brain volumes quoted in other strains of similar age and body weight.^7,42^

It is still possible that neuroanatomical differences exist between the Fischer 344 strain and other inbred (Lewis, Brown Norway) or outbred (Wistar, Sprague-Dawley, Long-Evans) strains that would cause co-registration between experimental scans to the Fischer atlas to fail. These validation studies have not yet been performed, so while it is highly likely that this atlas is relatively generalizable given the widespread use of other atlases in multiple rat strains, it should be used with care in strains other than the Fisher rat.

Some limitations of the present work are outlined below:

- The upsampled resolution of 60 μm isotropic may not allow delineation of very small structures. Additionally, adjacent grey matter nuclei, such as the thalamic nuclei and cortical nuclei, are very difficult to distinguish, and therefore are not delineated in this atlas.
- All rats used for this study were four months of age and are therefore considered to have reached mature adulthood. However, due to normal changes in brain shape and volume during the rat lifespan, this atlas may not be applicable to significantly older or younger rats.
- The sample sizes between male and female rats were uneven, with 24 male rats and 17 female rats. This could have the effect of over-weighting the structural differences that are observed in the male rats.
- Even though all of the rats used in this study were homozygous WT, they came from two different breeding schemes: 32 from homozygous WT male x WT female breeders and 9 from WT females x hemizygous Tg (TgF344-AD) male breeders. However, any effects of parental background are likely minimal, since a separate structural analysis found no significant anatomical differences between the homozygous wildtype rats bred from the WT/WT and WT/Tg parental backgrounds used in this study.
- Manual segmentation of all structures was performed in the coronal plane. Although the transverse and sagittal views were inspected to ensure spatial continuity of the various anatomical structures, it is impossible to guarantee smoothness of the structural boundaries in all three dimensions.

Ever increasing amounts of data are being generated in pre-clinical studies of rodent models, primarily due to the development of improved techniques to generate MR images. The Fischer 344 brain atlas presented in this paper can be used in preclinical studies to quickly and precisely identify anatomical regions and their volumes. The advantages of this tool are three-fold. First, this digital atlas allows researchers to expediently process large datasets by semi-automating the process of anatomical analysis and eliminating manual paper-based anatomical analysis. Second, this atlas reduces subject-wise bias in segmentation as all experimental subjects are mapped to the same set of labels. Third, this digital atlas can be used in combination with statistical software such as RMINC in the R environment to perform group-wise regions-of-interest comparisons. Our aim was to provide a novel tool for researchers working with Fischer 344 rats and thus we provide all files for open download at https://doi.org/10.5281/zenodo.3555556.

## Acknowledgements

This research was supported by the Canadian Institutes of Health Research (PJT - 148751, J.N.) and the Fonds de la recherche en santé du Québec (Chercheurs Boursiers # 35275, J.N.). G.A.D. is supported in part by funding provided by Brain Canada, in partnership with Health Canada, for the Canadian Open Neuroscience Platform initiative.

## Author Contributions Statement

D.G generated and manually segmented the template image. D.G, C.F, and J.N wrote the main manuscript text. C.F, D.M, and J.N collected data. J.G, M.C and GAD provided statistical analyses. All authors reviewed the manuscript.

## Additional Information

### Ethical approval

All applicable international, national, and/or institutional guidelines for the care and use of animals were followed. All animal procedures and experiments were approved by the McGill University Animal Care Committee (UACC).

### Data Availability Statement

The final anatomical segmentation volume along with the template brain, corresponding label descriptions, and a brain masking volume were each converted from MINC format into NIfTI format using the minc-toolkit `mnc2nii` function. All of the above are available for download and free use through the file repository Zenodo (DOI: 10.5281/zenodo.3555556) in both NIfTI and the MINC 2.0 format – compatible with all MINC tools, as well as Pydpiper and RMINC software

