## Supplemental Information for "An MRI-Derived Neuroanatomical Atlas of the Fischer 344 Rat Brain"

*Scientific Reports*

Dana Goerzen^a^, Caitlin Fowler^b^, Gabriel A. Devenyi^c,d^, Jurgen Germann^c^, Dan Madularu^c,d,e^, M. Mallar Chakravarty^b,c,d^, and Jamie Near^b,c,d^

a. Department of Neuroscience, McGill University, Montreal, Canada.

b. Department of Biological and Biomedical Engineering, McGill University, Montreal, Canada.

c. Centre d’Imagerie Cérébrale, Douglas Mental Health University Institute, Montreal, Canada.

d. Department of Psychiatry, McGill University, Montreal, Canada.

e. Centre for Translational Neuroimaging, Northeastern University, 02115, Boston, MA, USA

**Corresponding Author:**

Dana Goerzen

**Definition of individual brain structures**

**Ventricular System**

The ventricular system, which is made up of the lateral, 3^rd^, 3^rd^, and 4^th^ ventricle, and the aqueduct, was delineated on the basis of high intensity signal in the T_2_-weighted image. As the ventricular channels are continuous with CSF spaces surrounding the brain, arbitrary boundaries were placed at edge of the brain along the 3^rd^ ventricle, 4^th^ ventricle, and the cerebral aqueduct.

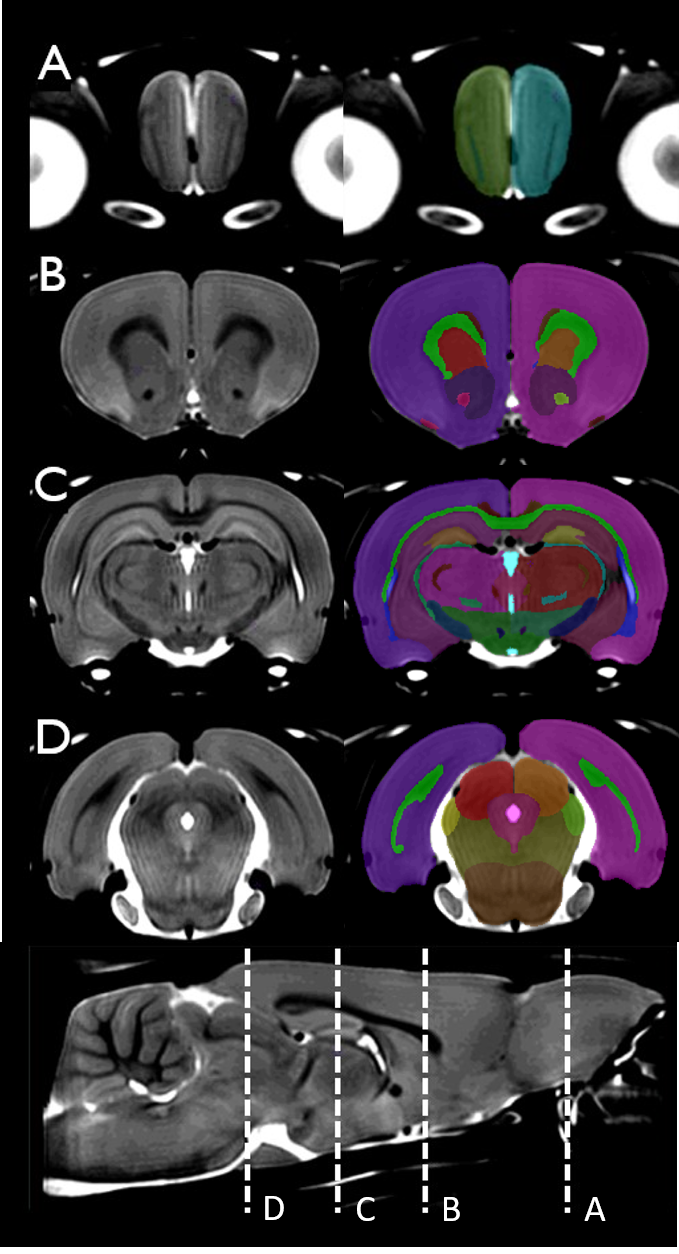

**Figure S1:** On the left column is the template brain, and on the right column is the template brain with the atlas file superimposed. This figure shows a series of coronal slices at different regions of the neocortex. Beginning rostrally in the Olfactory Bulb, at the rostrum of the Corpus Callosum, near the mid-Hippocampal Formation, and by the Superior Colliculus at the caudal neocortex, this figure serially shows the continuity of regions such as the Corpus Callosum.

**Hippocampal Formation**

The substructures in the hippocampal formation were delineated on the basis of contrast in the T_2_-weighted volume between the boundaries of CA3, CA2, CA1 subfields, which were delineated as one structure, and the dentate gyrus.

**Cerebellum**

As **Figure 2** indicates, the cerebellar lobules 1-10, Crus 1-2 Ansiform lobules, Paramedian lobe, Copula, and the Simple lobule were identifiable as unique parts of the cerebellum on the basis of high signal intensity of the T_2_-weighted image. They were further delineated into individual lobules by comparison of P&W atlas to the observed volume.

The arbor vitae/white matter was delineated as a single structure and was identifiable due to low signal intensity of white matter as compared to the surrounding cerebellar grey matter.

The flocculus and paraflocculus were identifiable as the flocculonodular node apart from the rest of the cerebellum. They were delineated apart from each other on the basis of a low signal intensity stripe separating the two lobes.

**Figure S2:** On the left is the template brain, and on the right is the template brain with the atlas file superimposed.Clear boundaries are identifiable between all of the major lobes. White matter and Arbor Vitae were indistinguishable based on the resolution of the image. All the hindbrain nuclei were delineated as one structure due to poor resolving power of the contrast levels in the hindbrain.
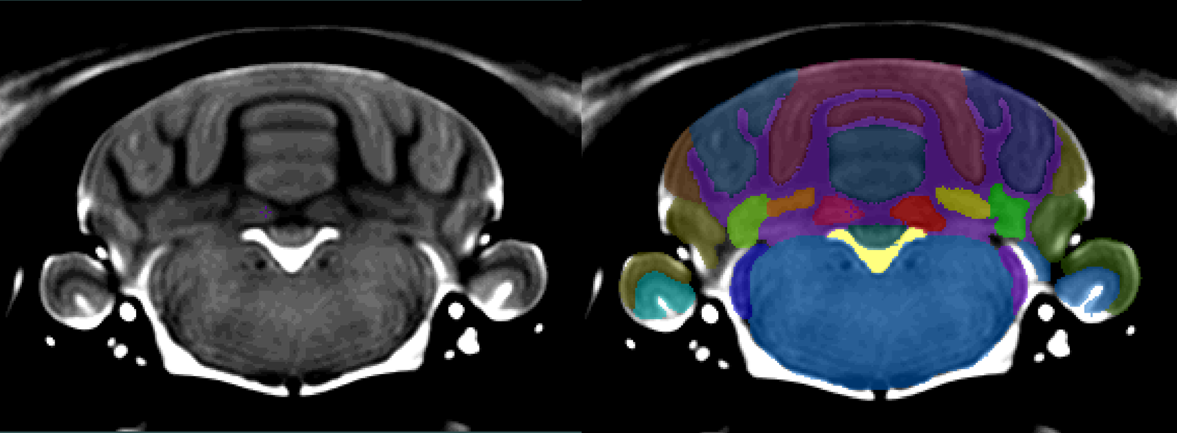

**Olfactory Bulb**

The olfactory bulb was segmented as a whole structure at the tip of the forebrain. The boundary between the brain and surrounding subarachnoid space has extreme contrast, with grey matter signal intensity being much greater. Olfactory bulb boundaries approaching the frontal cortex were identified on the basis of low signal intensity at the junction between the frontal cortex and the olfactory bulb.

**Hindbrain**

The hindbrain is made up of numerous nuclei and fibre tracts, such as the nucleus of the solitary tract, that were not identifiable in this segmentation. This region is distinct from the inferior cerebellum and the surrounding extra-brain space by signal intensity contrast, as well as from the surrounding extra-brain space. Boundaries on the rostral hindbrain were arbitrarily made at the rostral cerebellum. Anterior to the cerebellum is the inferior colliculus which is a midbrain structure. At the caudal hindbrain, the segmentation was ended where the hindbrain began to give way for the spinal cord.

**Pons**

The pons in this segmentation is made up of the pontine nuclei and surrounding rear-midbrain structures such as the raphe nuclei that were indistinguishable from each other on the basis of tissue contrast. Its caudal boundaries were identified as beginning at the caudal boundary of the neocortex. The pontine boundaries were differentiated from the inferior and superior colliculi as well as the neocortex and the caudal thalamus based on low signal intensity. Rostrally, the pons was differentiated from the midbrain using the cerebral peduncles low intensity fibre tract as a boundary.

**Midbrain**

Similarly to the hindbrain and pons, this structure contains several small nuclei that are indistinguishable on the basis of tissue contrast. This structure was segmented after many surrounding nuclei and structures that were segmented such as the pons, thalamus, corpus callosum and associated white matter, which allowed simple delineation along the body of this structure. Caudally, the midbrain structural boundary began at the cerebellar peduncles and extended rostrally until the main body of the thalamus.

### **Mammillary Bodies**

The mammillary bodies were bounded inferiorly by high intensity CSF. Caudally, the bodies ended and were distinct from the remaining body of the brain, and rostrally the bodies ended within the caudal hypothalamus. Superiorly, the mammillary bodies were delineated by comparing the low intensity region of the mammillary bodies to the higher intensity midbrain region.

**Basal Forebrain**

This region is made up of several nuclei and tracts that don’t present any identifiable tissue contrast given the current template. The basal forebrain was delineated after all surrounding identifiable nuclei and brain regions were segmented. Contains preoptic area, nucleus basalis, diagonal band of Broca, and substantia innominata.

**Bed Nucleus of the Stria Terminalis**

This structure was observed to have higher tissue intensity on the T_2_-weighted image compared to surrounding nuclei, with boundaries on one side being the low signal anterior commissure, posterior part, and on the other side the slightly lower signal thalamus. Segmentation was performed in conjunction with the P&W atlas.

**Periaqueductal Grey**

The periaqueductal grey is a concentric band of grey matter that surrounds the cerebral aqueduct. It was identified on the basis of higher signal contrast compared to the aqueduct and to the surrounding midbrain, superior colliculus, and inferior colliculus.

**Caudoputamen**

The caudoputamen was delineated after the corpus callosum was segmented due to the ease of corpus callosum (CC) segmentation. The inferior boundary was segmented based on the higher signal intensity of the caudoputamen compared to the nucleus accumbens.

### **Ventral Pallidum**

### The ventral pallidum was segmented in conjunction with the P&W atlas. It was distinctly lower in image contrast than the inferior neocortex, and was also lower in image contrast than the superior basal forebrain.

**Nucleus Accumbens**

The nucleus accumbens was delineated after the caudoputamen was segmented and showed lower signal intensity than surrounding tissues such as the olfactory tubercle and bed nucleus of stria terminalis as well as lower signal intensity than the neighbouring caudoputamen.

**Inferior Colliculus**

The caudal boundary of the inferior colliculus was segmented by contrast differences between the IC and the 4/5^th^ cerebellar lobules. The lateral boundaries of the inferior colliculus were segmented by the low signal intensity at the junction between IC and the neocortex. The rostral boundary of the inferior colliculus was delineated from the superior colliculus and pons on the basis of higher signal intensity of the inferior colliculus.

### **Superior Colliculus**

Superiorly, the superior colliculus was clearly bounded by the brain boundary. Moving caudally, the lateral boundary was increasingly bounded by the lateral brain boundary,. Medially, the superior colliculi were bordered by the periaqueductal grey, a high intensity region in the medial, caudal portion of the brain. The superior colliculi were inferiorly bounded by high voxel intensity midbrain.

### **Cochlear Nucleus**

The cochlear nucleus is found at the lateral boundary of the hindbrain, beneath the cerebellum. The lateral and superior boundaries based on contrast differences between surrounding CSF and the nucleus itself. The medial boundary was segmented by observing contrast differences between the hindbrain and the nucleus.

**Globus Pallidus**

The globus pallidus was segmented on the basis of lower signal intensity compared to the caudoputamen and higher signal intensity compared to the internal capsule and stria terminalis.

### **Septal Complex**

The septal nuclei were delineated on their lateral boundaries by the lateral ventricles. Superiorly, the septal complex was bounded by the low intensity corpus callosum and inferiorly the complex was bounded by the basal forebrain.

### **Median Preoptic Area**

The mdPOA was identifiable as being bounded on both sides by the medial septal complex which was distinctly lower in voxel intensity.

**Thalamus**

The thalamic nuclei were for the most part grouped together due to poor tissue contrast between individual nuclei within the region. The caudal boundary of the thalamic nuclei was identified based upon higher signal intensity of the T_2_-weighted volume than the midbrain. Moving rostrally, the thalamic nuclei is constrained by the corpus callosum and its associated white matter tracts. This boundary has very precise edge with low signal intensity white matter contrasted to the high signal thalamus. The inferior thalamic boundary was determined by differences in thalamic and hypothalamic signal intensity in conjunction with the P&W atlas in regions where the contrast was low. Rostral boundary was determined by the high signal bed nucleus stria terminalis. and 3^rd^ ventricle.

### **Subfornical Organ**

The small body of the subfornical organ was bounded laterally on both sides, rostrally and caudally, and inferiorly by the lateral ventricles. Superiorly, this structure was identifiably distinct from the fimbria due to contrasting image intensities.

**Hypothalamus**

The hypothalamic nuclei were grouped together as a single region due to poor signal contrast within the region. The superior boundary of the hypothalamus was delineated based upon tissue contrast between the hypothalamus and the thalamic nuclei. Its inferior boundary was segmented through high tissue contrast between hypothalamic grey matter and extra-brain space. The caudal hypothalamus was segmented by orienting the brain with the P&W atlas structures that were already delineated. The rostral hypothalamus was segmented by observing tissue contrast between the hypothalamus and forebrain. The lateral boundaries were delineated by low signal at the junction between hypothalamus and neocortex and high contrast between the external capsule and the hypothalamus.

**Substantia Nigra**

The substantia nigra was segmented as a single structure due to poor tissue contrast between pars reticulata and pars compacta. It was significantly lower in signal intensity than the medially located midbrain. Laterally, it was differentiable from the white matter cerebral peduncle by a slight contrast at the junction—corroborated by the P&W atlas.

**Fibre Tracts**

All of the fibre tracts were differentiated from the surrounding grey matter due to low signal intensity of white matter on the T_2_-weighted volume compared to grey matter.

The corpus callosum, external capsule, dorsal hippocampal commissure, and alveus were delineated as a single structure due to poorly defined boundaries on the T_2_-weighted image. The cingulum was the only associated white matter tract that had a defined boundary with the corpus callosum.

The anterior commissure was divided into the intrabulbar part, anterior part, and posterior part by consulting the P&W atlas and comparing morphology. These structures are all contiguous, and the line differentiating the intrabulbar part from the anterior part is fairly arbitrary.

The internal capsule was unambiguously segmentable except for near the caudal part, where it bordered the fimbria. This region was segmented by consulting the P&W atlas.

The optic chiasm and optic tract are contiguous at the rostral part and were segmented into two structures by consulting the P&W atlas.

The fornix, mammillothalamic tract, fasciculus retroflexus, lateral olfactory tract, stria medullaris thalamus, stria terminalis, external medullary lamina, commissure superior colliculus, commissure inferior colliculus, and posterior commissure were all easily identifiable through high contrast between surrounding grey matter and the fibre tract

The fimbria was bounded on the superior edge by the high signal hippocampal formation. There was a slight boundary at the lateral junction between the external capsule and the fimbria. The rostral, and inferior boundary was defined by the high signal contrast between the thalamic nuclei and the fimbria.

The cerebral peduncle was bounded laterally by high signal intensity of the neocortex, and medially by a weak boundary between the peduncle and the substantia nigra which was corroborated by the P&W atlas.

**White Matter of Cerebellum**

The white matter of cerebellum/ arbor vitae was segmented due to high tissue contrast between the molecular layers of the cerebellum and the white matter. It was not further subdivided into each sub region of the cerebellum due to poor tissue contrast within the white matter layers.

**Comparison of absolute volumes**

As described in **Section** **2.5 Statistical Analyses**, we performed a statistical comparison of the absolute volumes of each anatomical region between male and female rats. All regions were found to be significantly larger in males after FDR correction. The results are shown in Table S1 below.

| **Structure** | **Absolute Difference in Male-Female Volume (mm^3^)** | **P-Value** | **Significant After FDR** |
| --- | --- | --- | --- |
| Anterior Part of Anterior Commissure | 0.23 | <0.001 | Yes |
| Aqueduct | 0.38 | <0.001 | Yes |
| Basal Forebrain | 1.53 | <0.001 | Yes |
| Bed Nucleus of the Stria Terminalis | 0.47 | <0.001 | Yes |
| Caudoputamen | 5.23 | <0.001 | Yes |
| Cerebellar Lobule 1/2 | 1.77 | <0.01 | Yes |
| Cerebellar Lobule 10 | 0.37 | <0.001 | Yes |
| Cerebellar Lobule 3 | 1.15 | <0.001 | Yes |
| Cerebellar Lobule 4/5 | 3.18 | <0.001 | Yes |
| Cerebellar Lobule 6 | 1.47 | <0.001 | Yes |
| Cerebellar Lobule 7 | 0.43 | <0.001 | Yes |
| Cerebellar Lobule 8 | 0.52 | <0.001 | Yes |
| Cerebellar Lobule 9 | 1.37 | <0.001 | Yes |
| Cerebellar White Matter/Arbor Vitae of Cerebellum | 4.92 | <0.001 | Yes |
| Cerebral Peduncle | 0.54 | <0.001 | Yes |
| Cingulum | 0.74 | <0.001 | Yes |
| Cochlear Nucleus | 0.22 | <0.001 | Yes |
| Commissure of the Inferior Colliculus | 0.02 | <0.001 | Yes |
| Commissure of the Superior Collicuus | 0.04 | <0.001 | Yes |
| Copula | 0.53 | <0.001 | Yes |
| Corpus Callosum/Exernal Capsule | 6.01 | <0.001 | Yes |
| Cortex | 63.62 | <0.001 | Yes |
| Crus 1 Ansiform Lobule | 2.35 | <0.001 | Yes |
| Crus 2 Ansiform Lobule | 1.01 | <0.001 | Yes |
| Dentate Gyrus | 2.76 | <0.001 | Yes |
| Dentate Nucleus | 0.21 | <0.001 | Yes |
| Entopeduncular Nucleus | 0.07 | <0.001 | Yes |
| External Medullary Lamina | 0.16 | <0.001 | Yes |
| Fasciculus Retroflexus | 0.03 | <0.001 | Yes |
| Fastigial Nucleus | 0.24 | <0.001 | Yes |
| Fimbria | 0.76 | <0.001 | Yes |
| Flocculus | 1.10 | <0.001 | Yes |
| Fornix | 0.08 | <0.001 | Yes |
| Fourth Ventricle | 0.61 | <0.001 | Yes |
| Globus Pallidus | 0.60 | <0.001 | Yes |
| Hindbrain | 18.62 | <0.001 | Yes |
| Hippocampal Formation | 6.54 | <0.001 | Yes |
| Hypothalamus | 4.50 | <0.001 | Yes |
| Inferior Colliculus | 1.86 | <0.001 | Yes |
| Internal Capsule | 1.41 | <0.001 | Yes |
| Interposed Nucleus | 0.24 | <0.001 | Yes |
| Intrabulbar Part of Anterior Commissure | 0.12 | <0.001 | Yes |
| Lateral Olfactory Tract | 0.09 | <0.001 | Yes |
| Lateral Septum | 1.09 | <0.001 | Yes |
| Lateral Ventricle | 1.04 | <0.01 | Yes |
| Mammillary bodies | 0.18 | <0.001 | Yes |
| Mammillothalamic Tract | 0.03 | <0.001 | Yes |
| Medial Septum | 0.17 | <0.001 | Yes |
| Median Preoptic Nucleus | 0.01 | <0.001 | Yes |
| Midbrain | 7.22 | <0.001 | Yes |
| Nucleus Accumbens | 0.85 | <0.001 | Yes |
| Olfactory Nuclei | 14.59 | <0.001 | Yes |
| Optic Chiasm | 0.11 | <0.001 | Yes |
| Optic Tract | 0.79 | <0.001 | Yes |
| Paraflocculus | 1.49 | <0.001 | Yes |
| Paramedian Lobule | 1.22 | <0.001 | Yes |
| Periaqueductal Grey | 1.05 | <0.001 | Yes |
| Pons | 4.65 | <0.001 | Yes |
| Posterior Commissure | 0.03 | <0.001 | Yes |
| Posterior Part of Anterior Commissure | 0.07 | <0.001 | Yes |
| Simple Lobule | 2.11 | <0.001 | Yes |
| Stria Medullaris of the Thalamus | 0.06 | <0.001 | Yes |
| Stria Terminalis | 0.24 | <0.001 | Yes |
| Subfornical Organ | 0.02 | <0.001 | Yes |
| Substantia Nigra | 0.47 | <0.001 | Yes |
| Superior Colliculus | 2.56 | <0.001 | Yes |
| Superior Thalamic Radiation | 0.07 | <0.001 | Yes |
| Thalamus | 5.81 | <0.001 | Yes |
| Third Ventricle | 0.43 | <0.001 | Yes |
| Trochlear Nerve | 0.01 | <0.001 | Yes |
| Ventral Pallidum | 0.35 | <0.001 | Yes |

**Table S1.** Two sample, two tailed t-test comparing the absolute volumes for each structure between Male (n=24) and Female (n=17) subjects. Middle column displays the difference between Male Volume-Female Volume. As the right column shows, all structures were significantly larger in male brains, after Benjamini-Hochberg FDR correction.

**Voxel-wise DBM Variability Analysis**

In order to quantify the sensitivity of the DBM average at a voxel-wise level, we undertook a variability analysis of the final deformation field relative Jacobians using RMINC. We calculated the coefficient of variation (CV), voxel-wise for males and female groups, in order to quantify the variation across wildtype "genetically identical" subjects. Figures S3 and S4 show the CV across the brain for males and females respectively.

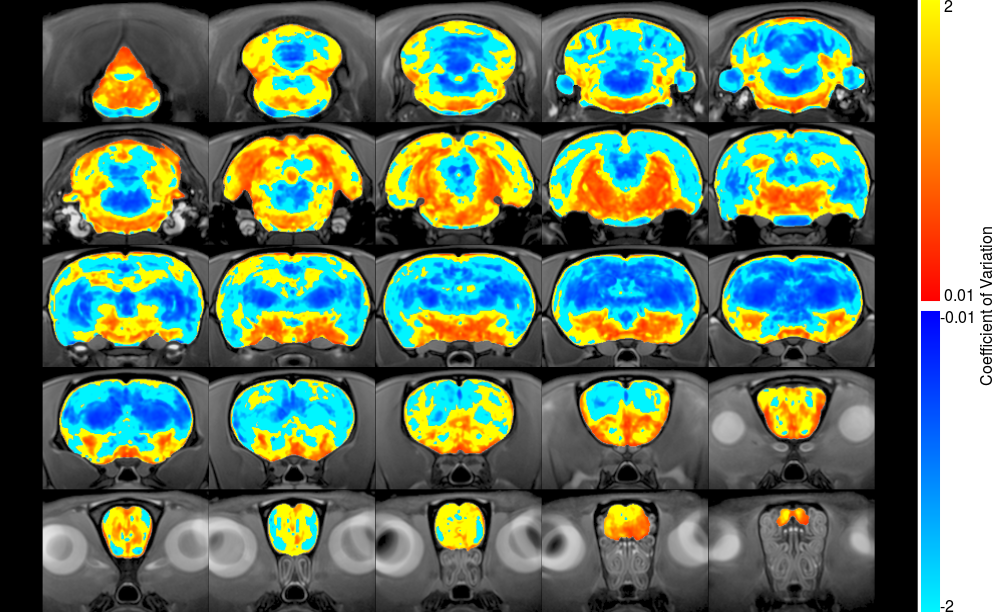

**Figure S3**: Voxel-wise map of male coefficient of variation.

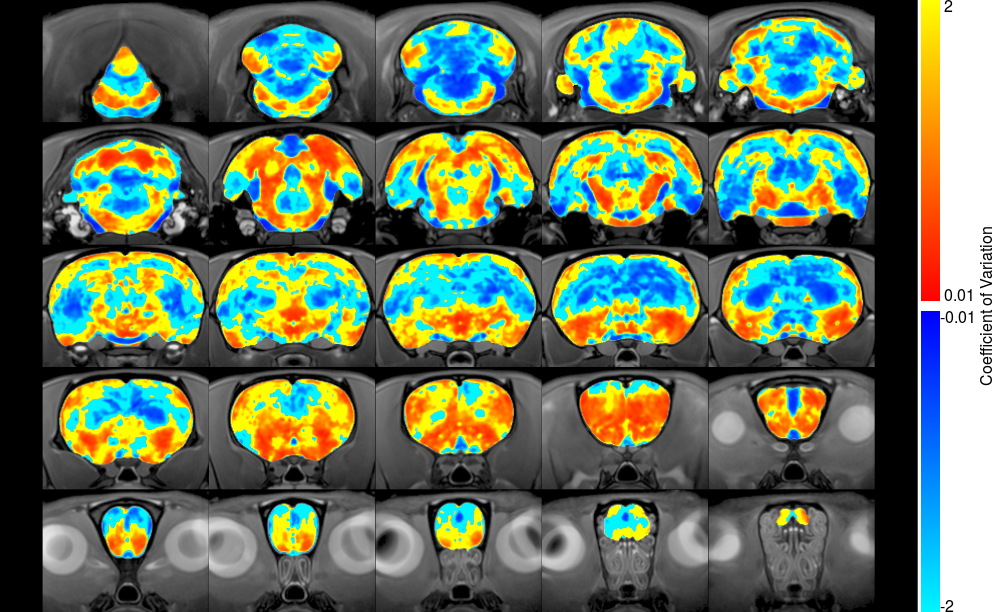

**Figure S4**: Voxel-wise map of female coefficient of variation.

**Tissue Probability Maps**

As described in Section **2.6 Tissue Probability Maps** of the main manuscript, ANTs Atropos was used to generate tissue probability maps of GM, WM, CSF, and other tissue based on the signal intensity histograms of the average brain template, along with the manually segmented tissue classes as a prior label image with a weight of 0.5. The resulting Classification image is shown in in Figure S5 below. The probability maps for each individual tissue type, as well as the unified classification map shown below are each included in the Zenodo repository along with the Fischer 344 template and atlas.

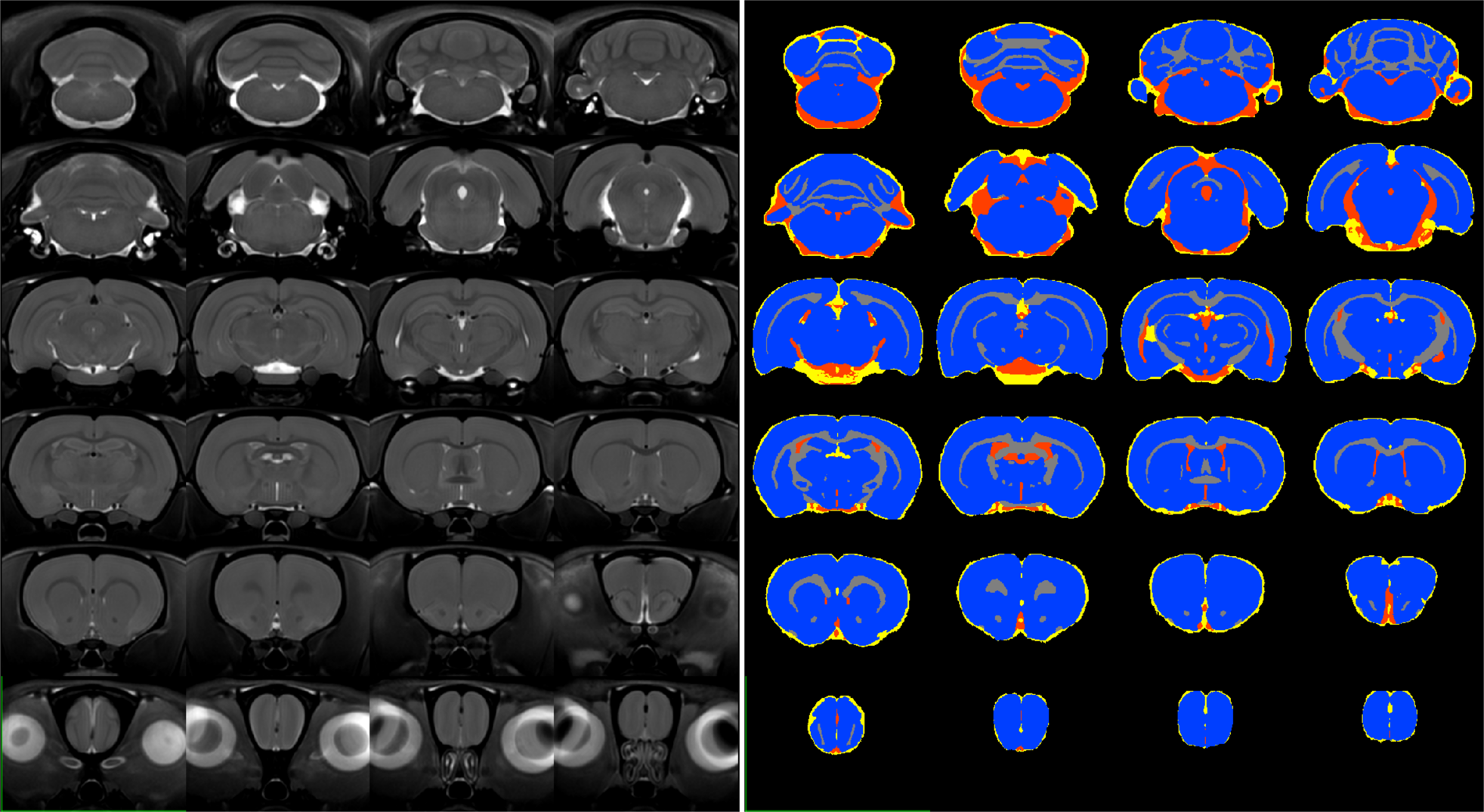

**Figure S5**. Tissue probability classification maps (right) generated using ANTs Atropos, and showing GM (blue), WM (grey), CSF (red) and other (yellow). The template from which these maps were generated is shown on the left.
